# Single-cell analysis of the ventricular-subventricular zone reveals signatures of dorsal and ventral adult neurogenic lineages

**DOI:** 10.1101/2021.02.10.430525

**Authors:** Stephanie A. Redmond, Arantxa Cebrian Silla, Marcos Assis Nascimento, Benjamin Mansky, David Wu, Kirsten Obernier, Ricardo Romero Rodriguez, Daniel A. Lim, Arturo Alvarez-Buylla

## Abstract

The ventricular-subventricular zone (V-SVZ) is home to the largest neurogenic niche in the adult mouse brain. Previous work has demonstrated that resident stem cells in different locations within the V-SVZ produce different subtypes of new neurons for the olfactory bulb. While great progress has been made in understanding the differences in regional stem cell potential using viral and genetic lineage tracing strategies, the core molecular heterogeneity that underlies these regional differences is largely unknown. Here we present single whole-cell and single nucleus sequencing datasets of microdissected adult mouse V-SVZ, and evidence for the existence of two broad populations of adult neural stem cells. By using spatially resolved microdissections in the single nucleus sequencing dataset as a reference, and mapping marker gene expression in the V-SVZ, we find that these two populations reside in largely non-overlapping domains in either the dorsal or ventral V-SVZ. Furthermore, we identified two subpopulations of newly born neurons that have gene expression consistent with dorsal or ventral origins. Finally, we identify genes expressed by both stem cells and the neurons they generate that specifically mark either the dorsal or ventral adult neurogenic lineage. These datasets, methods and findings will facilitate the study of region-specific regulation of adult neurogenesis.

## Introduction

Neural stem cells (NSC) persist in the adult mouse brain in the walls of the forebrain ventricles. This neurogenic niche includes the ventricular and subventricular zones in the walls of the lateral ventricles (V-SVZ), home to a subpopulation of astrocytes (B cells) that function as the NSCs (Chaker et al., 2016; Doetsch et al., 1997; Ihrie and Alvarez-Buylla, 2011; Lim and Alvarez-Buylla, 2014; Mirzadeh et al., 2008). B cells generate intermediate progenitors (C cells) that, in turn, give rise to neuroblasts (A cells) that migrate to the olfactory bulb (OB) (Obernier et al., 2018; Ponti et al., 2013). A subpopulation of B cells also generate oligodendrocytes (Figueres-Oriate et al., 2019; Gonzalez-Perez, 2014; Kazanis et al., 2017; Menn et al., 2006; Nait-Oumesmar et al., 1999; Picard-Riera et al., 2002). From the initial interpretation that adult NSCs are multipotent and able to generate a wide range of neuronal cell types (Morshead et al., 1994; Reynolds and Weiss, 1992; van der Kooy and Weiss, 2000), the field has moved to a model of adult NSCs being highly heterogeneous and specialized for the generation of specific types of neurons, and possibly glia (Chaker et al., 2016; Delgado et al., 2020; Fiorelli et al., 2015; Merkle et al., 2014, 2007; Tsai et al., 2012). Previous single-cell sequencing experiments in the V-SVZ have described the many broad classes of cells that reside in the niche.. For example, transcriptional analyses after cell sorting have identified stages in the B-C-A cell lineage (Borrett et al., 2020; Codega et al., 2014; Dulken et al., 2017; Xie et al., 2020), as well as populations of NSCs that appear to activate after injury (Llorens-Bobadilla et al., 2015). Profiling of the entire niche has highlighted differences between quiescent and activated B cells (Mizrak et al., 2020; Zywitza et al., 2018). However, the differences among B cells of equivalent activation state (e.g. quiescent, primed or activated) or the B cell heterogeneity that leads to the generation of diverse neuronal subtypes remain poorly understood.

NSC heterogeneity, interestingly, is largely driven by their location within the adult V-SVZ. This concept explains why young neurons in the OB originate over such a wide territory in the walls of the lateral ventricles. Multiple studies have begun to identify regional differences in gene expression among the lateral, septal and subcallosal walls of the lateral ventricles. For example, differences in gene expression of B cells from the septal and lateral walls of the lateral ventricles have been recently observed (Mizrak et al., 2019). Other studies have shown that *Pax6* and *Hopx*-expressing cells correspond to dorsal V-SVZ progenitors (Hack et al., 2005; Kohwi et al., 2005; Zweifel et al., 2018), and *Vax1*-expressing young neuroblasts are derived from ventral progenitors (Core et al., 2020). Spatially defined lineage tracing studies using microinjections of viruses have identified subdomains of the V-SVZ that give rise to specific subtypes of OB neurons (Merkle et al., 2014, 2007; Ventura and Goldman, 2007). Lineage tracing studies have demonstrated that these regional subdomains largely follow the territories defined by developmentally regulated transcription factors including Pax6, Nkx2.1, Nkx6.2, and Emx1 (Delgado et al., 2020; Delgado and Lim, 2015; Kohwi et al., 2007, 2005; Merkle et al., 2014; Willaime-Morawek et al., 2006; Young et al., 2007), but the molecular differences among B cells underlying their regionally-restricted potential are largely unknown.

Here we have undertaken single-cell and single-nucleus RNA sequencing of the microdissected V-SVZ to gain insight into these important questions regarding NSC heterogeneity and their developmental potential. Clustering analysis reveals strong dorso-ventral differences in lateral wall B cells. Validation of these differential gene expression patterns has revealed the anatomical boundary that separates these dorsal and ventral B cell domains. Additionally, our analysis identifies subpopulations of A cells defined by maturation state and dorso-ventral origin. We also find that a subset of dorso-ventral B cell transcriptional differences are retained through the C and A cell stages of the lineage. These new data advance our molecular understanding of how major region-specific neural lineages emerge in the adult V-SVZ and begin to delineate major functional subclasses of adult-born young neurons.

## Results

### Single-cell RNA sequencing distinguishes B cells from parenchymal astrocytes and reveals B cell heterogeneity

For whole single-cell RNA sequencing (scRNA-Seq), we microdissected the lateral wall of the lateral ventricle from hGFAP:GFP mice at postnatal day (P) 29-35 (n=8, Fig 1A; Fig 1-S1A). To determine possible sex differences in downstream analyses, two male and two female samples (n=4 samples total) were dissociated and multiplexed by labeling cells with sample-specific MULTI-seq barcodes (McGinnis et al., 2019). Multiplexed samples were then pooled for the remainder of the single-cell isolation protocol. Two technical replicates of pooled samples were loaded in separate wells of the Chromium Controller chip (10x Genomics) for single-cell barcoding and downstream mRNA library preparation and sequencing (Fig 1A). Cells carrying multiple barcodes or a high number of mRNA reads (4,128 out of 35,025 cells, 11.7%) were considered doublets and were eliminated from downstream analysis. Data from each technical replicate were integrated for batch-correction (Stuart et al., 2019). We then performed unbiased clustering of cell profiles and calculated UMAP coordinates for data visualization (Fig 1A). The clustering of lateral wall V-SVZ cells was not driven by sample, technical replicate, or sex (Fig 1-S1B-J). Cell cluster identities were annotated based on the detection of known cell type markers (Fig 1B-C). We identified 37 clusters overall, with 14 clusters corresponding to cell types within the neurogenic lineage: NSCs (B cells), intermediate progenitors (C cells), and neuroblasts (A cells) (Doetsch et al., 1999; Obernier et al., 2018). In addition, our analysis identified cell clusters corresponding to parenchymal astrocytes, ependymal cells, neurons, oligodendroglia, microglia, pericytes, vascular smooth muscle cells, and endothelial cells (Fig 1B-C).

**Figure 1.**
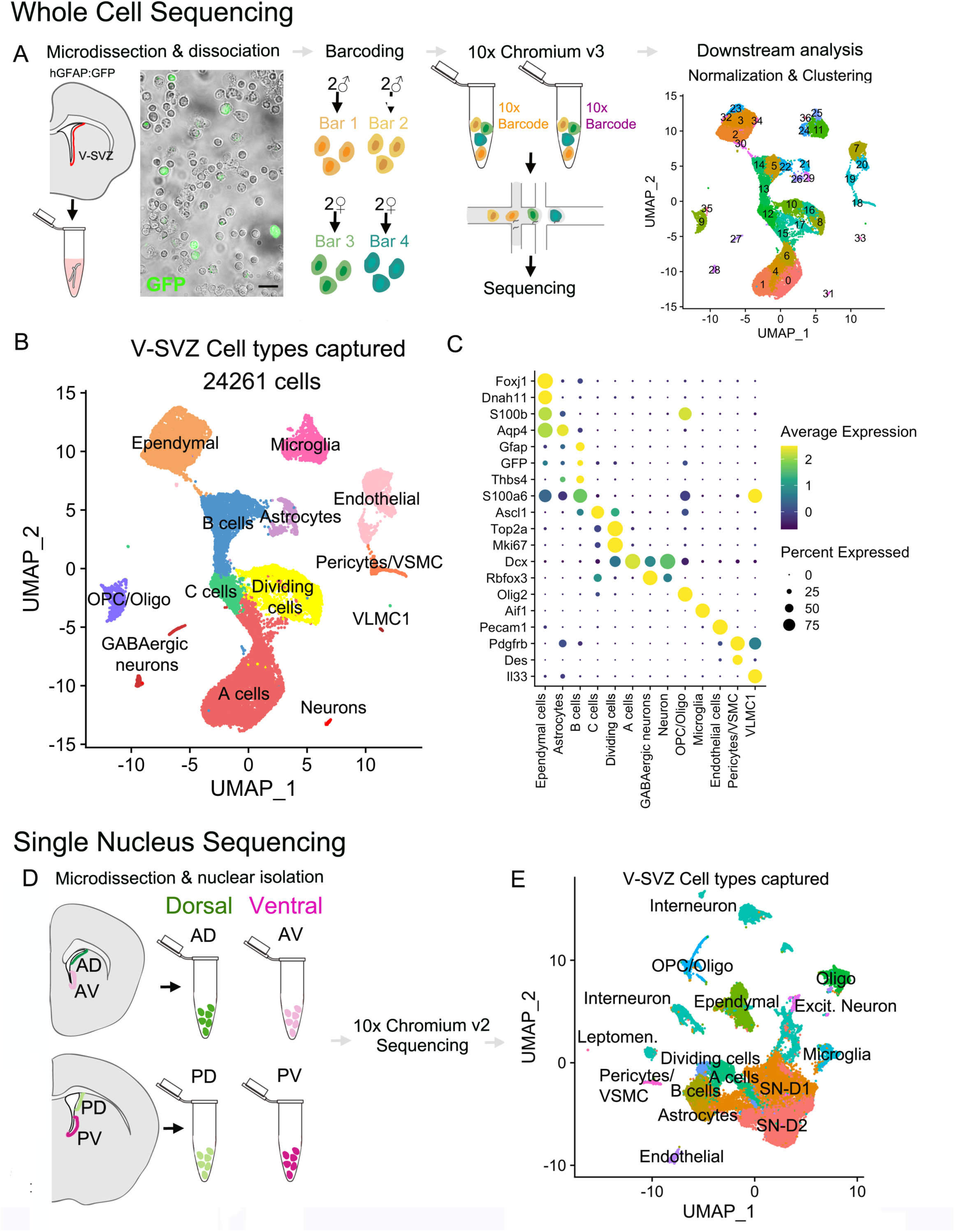
Whole-cell and single nucleus sequencing captures the cellular diversity & activation cascade of the adult neurogenic niche. A. Schematic of the whole-cell single cell isolation and sequencing protocol (scRNA-Seq). The lateral wall of the lateral ventricles was microdissected from young adult hGFAP:GFP mouse brains. Four samples were multiplexed with MULTI-seq barcodes and combined together. Two 10x Chromium Controller lanes were loaded as technical replicates, and cells were sequenced and processed for downstream analysis. B. UMAP plot of scRNA-Seq cell types after demultiplexing and doublet removal. C. Dot plot of cell type specific marker expression in the clusters from (B). D. Schematic of single nucleus sequencing (sNucRNA-Seq). Anterior ventral (AV), anterior dorsal (AD), posterior ventral (PV) and posterior dorsal (PD) regions were microdissected from young adult CD1 mice. Each region was processed for sNucRNA-Seq in parallel and loaded into its own 10x Chromium Controller lane. E. UMAP plot of sNucRNA-seq cell types identified from all four regions after quality control processing steps.

NSCs in the V-SVZ correspond to a subpopulation of astrocytes (B cells) derived from radial glia (Doetsch et al., 1999; Laywell et al., 2000; Merkle et al., 2004). B cells not only have astrocytic ultrastructure but also express many astrocytic markers (Borrett et al., 2020; Codega et al., 2014). Therefore, identifying markers that distinguish parenchymal astrocytes from B cells has been a challenge in the field. In our dataset, B cell and striatal astrocyte clusters are both defined by *Gfap* and *GFP* expression (Fig 1-S2A-B), among other genes expressed in common. We classified striatal astrocyte clusters based on increased expression of *Aqp4, S100b, and Cxcl14* (Codega et al., 2014; Zywitza et al., 2018), while B cell clusters were defined by expression of *S100a6* (Kjell et al., 2020) (Fig 1-S2A-B). Using these classifications, we performed differential gene expression analysis between B cells and astrocytes (clusters 5, 14, and 22 vs. clusters 21, 26, and 29, respectively). We found that B cells had a higher expression of *Maff, Zfp36, Bex4, Lgals3, and Anxa2* compared to striatal astrocytes, which in turn were enriched for *Clmn, Atp13a4, Eps8, Pcdh7, and Syne1* (Fig 1-S2C, Supplementary Table 1).

To better understand biological differences between striatal astrocytes and B cells, we performed gene ontology (GO) enrichment on the differentially expressed genes. Genes associated with synapse regulation (GO:0051965 and 0051963), macroautophagy (GO: 0016241), and dendrite development and morphogenesis (GO: 0016358 and 0048813), among others, were overrepresented in striatal astrocytes compared to B cells (Fig 1-S2D). In contrast, B cells were enriched in terms associated with RNA regulation (GO:0000463, 0034471, 0000956, and 0000966) and mitochondrial regulation (GO:006626, 0073655, 0090201, and 0010823) (Fig 1-S2D). These differentially represented GO terms support the known neuro-regulatory function of astrocytes, as well as the increased transcriptional regulation that has been associated with the transition of NSCs from quiescent to activated states (Dulken et al., 2017; Llorens-Bobadilla et al., 2015). Using this cell type classification, we included B cell clusters, but not striatal astrocytes, in our downstream analysis of the neurogenic lineage.

### scRNA-Seq captures neurogenic progression in the V-SVZ

In our scRNA-Seq dataset, we found that the majority of cells were part of the neurogenic lineage, which is composed of primary progenitors, intermediate progenitors, and young neurons (the B-C-A cell lineage). These neurogenic clusters were in the center of the UMAP plot. With this analysis, the data had a “hummingbird-like” shape with B cells (clusters 5, 13, 14 and 22) at the top in the bird’s head and neck, proliferating cells (clusters 8, 10, 16, and 17) and C cells (cluster 12) resembled the bird’s body and wing, and A cells (clusters 0, 1, 4, 6 and 15) formed a tail (Fig 2A). As expected, all B cell clusters expressed *Gfap, GFP,* and *S100a6* (Fig 2B-D). We identified a subpopulation of B cells as the quiescent B cells (Codega et al., 2014; Llorens-Bobadilla et al., 2015) (clusters 5, 14 and 22) characterized by high expression of *Thbs4* and *Gfap,* no *Egfr* and low *Ascl1* expression (Fig 2E, B, F-G). In contrast, cluster 13 (the neck region of the “hummingbird”) corresponded to activated B cells with lower expression of *Gfap, GFP,* and *S100a6,* but high *Egfr* and *Ascl1* expression (Codega et al., 2014) (Fig 2B-D, F-G). Cluster 13 cells also expressed *Notum*, a marker recently associated with activating qNSCs (Mizrak et al., 2020) (Fig 2H). Bordering cluster 13 in the chest region of the ‘bird’ was cluster 12, identified as the C cell cluster, which had low expression of astrocytic markers *Gfap* and *GFP* (Fig 2B-C), but high expression of *Egfr* and *Ascl1* (Fig 2F-G). The wing area of the ‘bird’ contained *mKi67*+ proliferating cells (Fig 2I), and in the tail area of the ‘bird’ below, clusters corresponding to neuroblasts were characterized by high expression of Dcx+ (Fig 2J). We used cell-cycle scoring to classify cells by their G2M, S, or G1 phase (Tirosh et al., 2016), and found that a high number of cells in cluster 12 were in the S phase (691/992, 69.6%), consistent with cluster 12 corresponding to intermediate progenitors (C cells) (Fig 2K).

**Figure 2.**
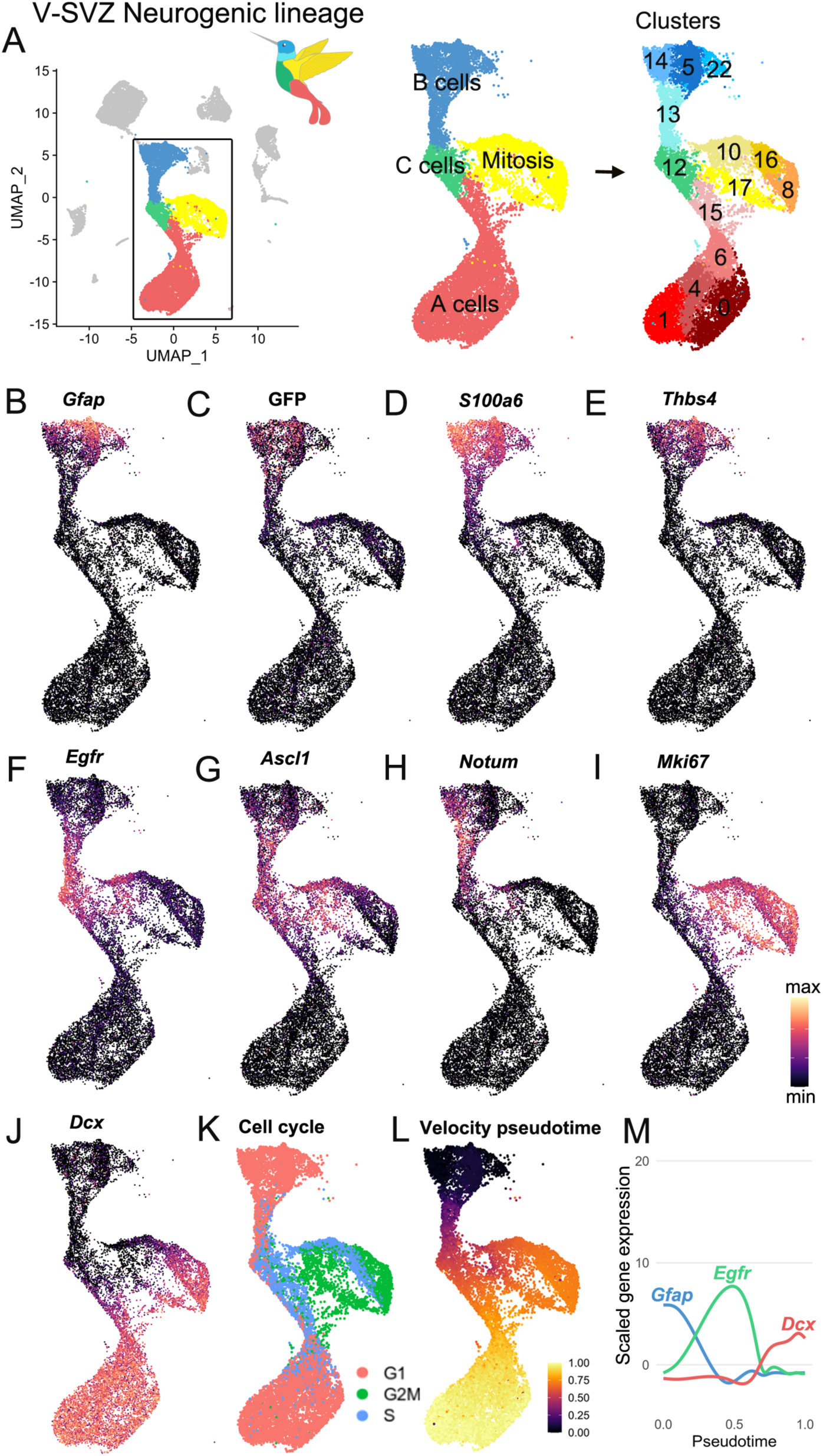
Characterization of the scRNA-Seq V-SVZ neurogenic lineage A. The neurogenic lineage has a “bird-like” shape, with B cells forming the head (blue), C cells in the body (green), dividing cells in the wing (yellow), and A cells in the tail (red). These are divisible into 14 distinct clusters, including 4 B cell clusters, 1 C cell cluster, 4 clusters of dividing cells, and 5 A cell clusters (right). B. -J. Gene expression captures progression along the lineage, with canonical markers of each stage expressed in its corresponding region of the UMAP plot. K. Scoring cells by phase of the cell cycle reveals cells in G2M and S phase occupying the wing of the bird. L. Pseudotime calculated by RNA velocity recapitulates the B to C to A trajectory along the neurogenic lineage. M. Genes associated with B cells, activated B and C cells, and A cells peak in expression at corresponding stages in the lineage.

The unbiased clustering analysis pooled dividing cells into the wing region of the ‘hummingbird’. A closer look at these clusters of mitotic cells in G2 or metaphase (G2M; clusters 10, 16, 17, and 8) showed that their gene expression pattern overlaps with that of the of non-dividing neurogenic lineage cell progression: Cluster 10 expresses markers of B cells (*GFP, Notum*), cluster 16 had markers of C cells (*Ascl1*), and cluster 8 expresses markers of A cells (*Dcx*) (Fig 2B-K). This suggests that these clusters correspond to dividing B, C, and A cells, respectively, which have all been previously observed by electron and confocal microscopy *in vivo* (Doetsch et al., 1997).

Identified by *Dcx* expression, the largest number of cells within the neurogenic lineage corresponded to neuroblasts and young neurons (A cells) (clusters 0, 1, 4, 6 and 15; the tail region of the hummingbird) (Fig 2A, J). Interestingly, unbiased clustering subdivided A cells into 5 subclusters with different gene expression profiles. Consistent with previous work showing that a subpopulation of newly generated neurons continues to divide (Lois and Alvarez-Buylla, 1993; Menezes et al., 1995), clusters 6 and 15 contain *Dcx*+ A cells with proliferative markers (e.g. *mKi67*) (Fig 2I-K). Genes that distinguished clusters 0, 1, 4, 6 and 15 are discussed below. The overall progression from B-C-A cells described above is supported by RNA-velocity lineage trajectory reconstruction (La Manno et al., 2018), in which genes defining B, C and A cells are expressed sequentially in distinct phases in pseudotime (Fig 2L-M).

Overall, our single-cell dataset recapitulates the known B-C-A cell progression through the neurogenic lineage. Intriguingly, our analysis also reveals heterogeneity among B, C, and A cells, in which each cell type is subdivided into multiple distinct clusters. Among C cells, the heterogeneity was mostly driven by different stages of the cell cycle (Fig 2F-G, I, K). What drives heterogeneity among B and A cell clusters?

### Single nucleus sequencing (sNucRNA-Seq) from microdissected V-SVZ regions captures V-SVZ and striatal cell types

To complement our single-cell transcriptomic analysis, we performed single-nucleus sequencing (sNucRNA-Seq) from microdissected V-SVZ subregions of P35 CD1 mice (n=8 males, 9 females). We isolated single nuclei from four distinct microdissected quadrants of the V-SVZ: the anterior-dorsal (AD), posterior-dorsal (PD), anterior-ventral (AV), and posterior-ventral (PV) regions (Fig 1D; Fig 1-S3A) (Mirzadeh et al., 2008). The four region samples (AD, PD, AV, and PV) were then processed in parallel for sNucRNA-Seq (Fig 1D; Fig 1-S3B). Our sNucRNA-Seq dataset contains 45,820 nucleus profiles. The four region samples underwent quality control steps of filtering out low-quality cells and putative doublets (see Methods). Data from each sample were combined and integrated (Seurat v3 *IntegrateData*) (Stuart et al., 2019), then clustered as the scRNA-Seq dataset above. Cell identities were annotated based on the detection previously described cell type markers (Martin et al., 2019; Zeisel et al., 2018) (Fig 1E; Fig 1-S3C-E).

To understand the differences between the scRNA-Seq and sNucRNA-Seq datasets, we compared the numbers of unique molecular identifiers (UMIs) and genes identified per cell. The scRNA-Seq cells have 3-fold more UMIs/cell (scRNA-Seq: 9,181; sNucRNA-Seq: 3,062 median UMIs/cell) and 2.1-fold more genes identified per cell (scRNA-Seq: 3,549; sNucRNA-Seq: 1,679 median genes/cell) (Fig 1-S3I). The two datasets also differed in the proportion of V-SVZ cell types represented (Fig 1-S3J).

In the sNucRNA-Seq data, we identified 42 clusters, including those corresponding to cell types within the neurogenic lineage: NSCs (B cells), mitotic intermediate progenitors (C cells), and neuroblasts (A cells) (Fig 1E, Fig 1-S3C-E). Based on the B cell- or astrocyte-specific markers identified in the scRNA-Seq data above, we also identify a parenchymal astrocyte cluster, as well as ependymal cells, striatal neurons, oligodendroglia, microglia, pericytes and vascular smooth muscle cells, endothelial cells, and leptomeningeal cells (Fig 1E, Fig 1-S3D-E). We also found that all four regions contributed to most clusters (Fig 1-S3F-H).

Given the greater proportion of high-quality neurogenic lineage cells, we focused the following analysis on the scRNA-seq dataset, making key supporting observations using the sNucRNA-Seq dataset.

### Quiescent B cell clusters correspond to regionally organized dorsal and ventral domains

As shown above, in our scRNA-Seq dataset, we found that quiescent B cells (the head of the ‘bird’; Fig 2A) were subdivided into three clusters: B cell cluster 5 (B(5)), B(14), and B(22) (Fig 3-S1A). To understand their molecular differences, we conducted differential expression analysis to identify significantly upregulated genes in each of the three B cell clusters (Fig 3-S1Bi) and candidate cluster-specific marker genes (Fig 3-S1Bii). When we examined the top ten candidate markers for each cluster, we found genes corresponding to known markers of dorsal and ventral B cell identity (Fig 3-S1C). For example, *Nkx6.2*, a transcription factor expressed in the ventral embryonic and postnatal ventricular zone (Merkle et al., 2014; Moreno-Bravo et al., 2010), is enriched in cluster B(14), as are *Notum* and *Lmo1* (Fig 3A, Fig 3-S1C) (Borrett et al., 2020; Mizrak et al., 2020). Similarly, *Gsx2*, a marker of dorsal B cells, is a marker of cluster B(5) (Fig 3B, Fig 3-S1C). Another known dorsal marker, *Emx1*, is significantly upregulated in cluster B(22) (Fig 3C, Supplementary Table 2). When we overlay the expression of these cluster markers on the neurogenic lineage UMAP plot, we find that their expression is largely restricted to cells within a single B cell cluster, and in the case of *Gsx2*, is retained in C cells and early-stage A cells (Fig 3A-C). Activated B cells in cluster B(13) also showed the expression of these regional markers but did not separate into multiple clusters at this resolution (Fig 2A). The distinct regional signature that drives clustering for quiescent B cells seems to be partially lost as these primary progenitors move into the activated state.

**Figure 3.**
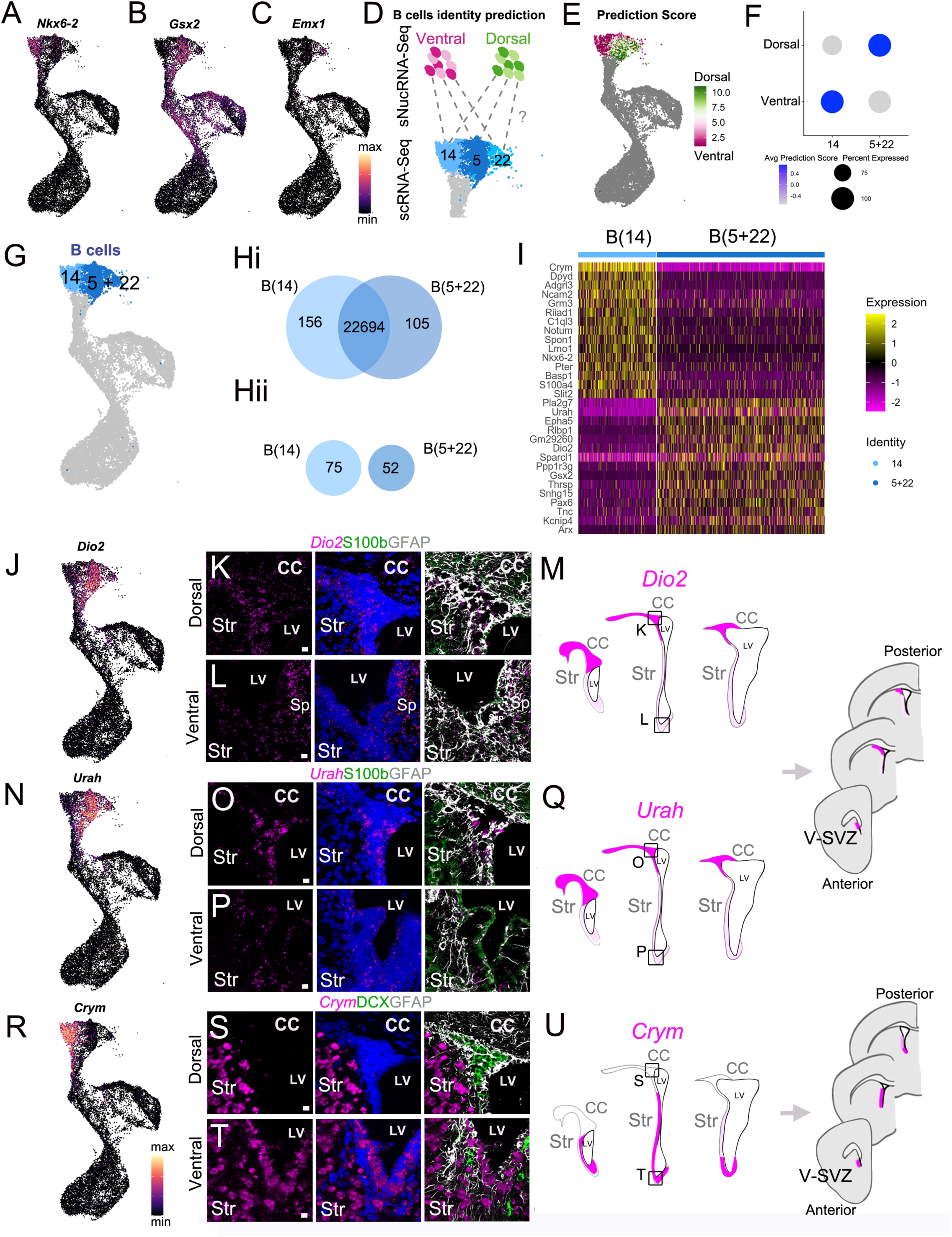
scRNAseq reveals regional heterogeneity among adult neural stem cells A. -C. UMAP plots of *Nkx2.1* (A), *Gsx2* (B), and *Emx1* (C) expression in the scRNA-Seq neurogenic lineage. D. Schematic of the region identity prediction calculation, where anchor gene sets (dashed gray lines) are calculated between Ventral (magentas) and Dorsal (greens) sNucRNA-Seq B cell nuclei and scRNAseq B cells (blues) and each scRNAseq B cell is given a Dorsal or Ventral predicted identity score. E. The net predicted identity scores of each scRNA-Seq B cell plotted in UMAP space, where strongly Dorsal predicted identity cells are dark green, and strongly Ventral predicted identities are dark magenta. Dark gray cells were not included in the analysis. F. Dot plot of the average Dorsal or Ventral predicted identity scores for scRNA-Seq B cell clusters B(14) and B(5+22). G. UMAP plot of B cell cluster identities used in the following analysis: B(14) (light blue) and B(5+22) (dark blue). H. ***i.*** Venn diagram summarizing differential gene expression analysis between clusters B(14) (light blue) and B(5+22) (dark blue). ***ii.*** Numbers of candidate marker genes identified after selecting significantly upregulated genes expressed in no more than 40% of cells of the other cluster. I. Heatmap depicting expression of the top 10 differentially expressed genes between clusters B(14) (left) and B(5+22) (right). J. UMAP plot of *Dio2* expression in the scRNA-Seq neurogenic lineage. K. - L. Confocal micrograph of *Dio2* RNA (magenta), S100b (green) and GFAP (white) protein expression in the dorsal (K) and ventral (L) V-SVZ. M. Summary schematic of anterior to posterior V-SVZs of each section (left) and coronal brain sections (right) denotes strong (magenta), sparse (light magenta) or absent (white) *Dio2* expression. Boxed areas in the center V-SVZ schematic denote locations of dorsal (K) and ventral (L) high magnification images. N. UMAP plot of *Urah* expression in the scRNA-Seq neurogenic lineage. O. - P. Confocal micrograph of *Urah* RNA (magenta), S100b (green) and GFAP (white) protein expression in the dorsal (O) and ventral (P) V-SVZ. Q. Summary schematic of anterior to posterior V-SVZs of each section denotes strong (magenta), sparse (light magenta) or absent (white) *Urah* expression. Boxed areas in the center V-SVZ schematic denote locations of dorsal (O) and ventral (P) high magnification images. R. UMAP plot of *Crym* expression in the scRNA-Seq neurogenic lineage. S. - T. Confocal micrograph of *Crym* RNA (magenta), DCX (green) and GFAP (white) protein expression in the dorsal (S) and ventral (T) V-SVZ. U. Summary schematic of anterior to posterior V-SVZs of each section (left) and coronal brain sections (right) denotes strong (magenta), sparse (light magenta) or absent (white) *Crym* expression. Boxed areas in the center V-SVZ schematic denote locations of dorsal (S) and ventral (T) high magnification images. DAPI: blue, LV: lateral ventricle, CC: corpus callosum, Str: striatum. Scale bars: 10 µm (K, L, O, P, S and T).

To test the hypothesis that B cells are dorso-ventrally organized in the V-SVZ, we took advantage of the region-specific microdissection of the sNucRNA-Seq cells (Fig 1D-E; Fig 1-S3) and metadata Label Transfer to predict scRNA-Seq B cell region identity (Stuart et al., 2019). Each B cell was assigned both a dorsal and ventral ‘predicted identity’ score based on their similarity to dorsal and ventral nuclei (Fig 3D). We then calculated the difference between dorsal and ventral scores for each scRNA-Seq B cell. We found that cells within each cluster were strongly dorsal-scoring (green) or ventral-scoring (magenta), with relatively few cells having similar dorsal and ventral prediction scores (gray) (Fig 3E). We found that cluster B(14) scored more highly for ventral identity on average, while clusters B(5) and B(22) scored more highly for dorsal identity (Fig 3-S1D).

To investigate the potential dorso-ventral spatial organization of dorsal-scoring clusters B(5) and B(22) *in vivo*, we performed RNAscope in situ hybridization for the differentially expressed transcripts *Small Nucleolar RNA Host Gene 15* (*Snhg15*, a lncRNA), and *Contactin-Associated Protein 2* (*Cntnap2*), which are upregulated in clusters B(5) and B(22), respectively (Fig 3-S1E-L). Overall, we found that *Snhg15* and *Cntnap2* probes did not exclusively localize in B cells, but were also expressed in subsets of A cells, C cells, and ependymal cell layer, making it difficult to confirm their B cell expression along the dorsal-ventral axis of the V-SVZ (Fig 3-S1F-H, J-L). To identify markers more specific to B cells within the V-SVZ, and test the hypothesis that B(5) and B(22) both correspond to dorsally-localized B cells, we combined them into a single cluster, B(5+22). Looking at this new cluster’s predicted dorsal and ventral identity scores, we found that it had a much higher average dorsal predicted identity score, as well as lower average ventral predicted identity score than when scored separately, nearly the inverse of cluster B(14)’s prediction scores (Fig 3F). This unsupervised, unbiased prediction of region identity at the single-cell level, based on sNucRNA-Seq region-specific microdissection, reinforces our observation that scRNA-Seq B cell clusters have strong gene expression signatures of dorsal or ventral V-SVZ identity.

We then asked what genes were differentially expressed between the putative dorsal cluster B(5+22) and the putative ventral cluster B(14) (Fig 3G-I). Among the differentially expressed genes were other known markers of dorsal B cells, such as *Pax6 and Hopx* (Fig 3 Hi, Supplementary Table 2), as well as novel dorsal-domain marker candidates such as *Urah* and *Dio2* (Fig 3Hii, I). We found that *Urah* and *Dio2* were highly expressed among GFAP+ cells in the V-SVZ dorsal “wedge” (the dorso-lateral corner of the V-SVZ enriched in B, C, and A cells), while lower expression was observed in the intermediate and ventral regions (Fig 3J-Q). This pattern of expression was conserved from anterior to posterior sections (Fig 3M, Q). To determine if the dorsal *Urah* and *Dio2* domain correspond to the dorsal *Hopx+* B cell population (Zweifel et al., 2018), we performed RNAscope *in situ* hybridization for *Hopx*. *Hopx* was expressed in B cells in the dorsal wedge with no expression in the intermediate and ventral V-SVZ regions (Fig 3-S1M-Q). Interestingly, low and high *Hopx*+ cells were observed in the wedge*. Hopx^high^* cells showed a sub-callosal localization forming a band that extended laterally from the septal corner of the V-SVZ, where Hopx+ cells have been previously described (Zweifel et al., 2018) (Fig 3-S1N-O, Q). We confirmed that HOPX protein followed the expression pattern of its mRNA by immunostaining (Fig 3-S1R).

Cluster B(14) marker *Crym* was expressed in intermediate and ventral V-SVZ GFAP+ cells (Fig 3R-U), with no expression in the V-SVZ dorsal wedge (Fig 3S). Interestingly, the *Crym*+ domain decreased in size along the anterior-posterior axis (Fig 3U). Consistent with our single-cell data, A cells identified by DCX expression were *Crym*-negative (Fig 3R-T). We also found that cells in the striatum expressed *Crym,* as previously described (Fig 3S-T) (Chai et al., 2017; Mizrak et al., 2019). To determine if *Crym* was expressed at the protein level in a regional pattern, we analyzed CRYM expression in P28 hGFAP:GFP mice. Consistent with the RNAscope assay, ventral SVZ GFP+ B cells were CRYM+, while dorsal wedge GFP+ cells were negative (Fig 4A-C). To better understand the antero-posterior distribution of the CRYM+ ventral domain, we analyzed B1 cell CRYM expression in V-SVZ whole mounts (Mirzadeh et al., 2008). Consistent with *Crym* expression patterns in coronal sections along the anterior-posterior axis (Fig 3U), we found the CRYM+ B cell domain in the wholemount covered ventral and intermediate domains of the lateral wall, but was absent from the dorsal and wedge area (Fig 4D). This wholemount analysis also shows how the CRYM+ domain decreases in size and becomes restricted to more ventral regions along the anterior to posterior axis (Fig 4D-G). Using the hGFAP-GFP mice and staining with BETA-CATENIN to reveal pinwheels and apical contacts of B1 cells, we confirmed that CRYM was expressed in GFP+ ventral B1 cells, while absent in dorsal B1 cells (Figure 4H-I). Taken together, we conclude that *Crym* transcript and protein expression defines a wide ventral territory that decreases caudally, and is absent from the dorsal and wedge V-SVZ regions.

**Figure 4.**
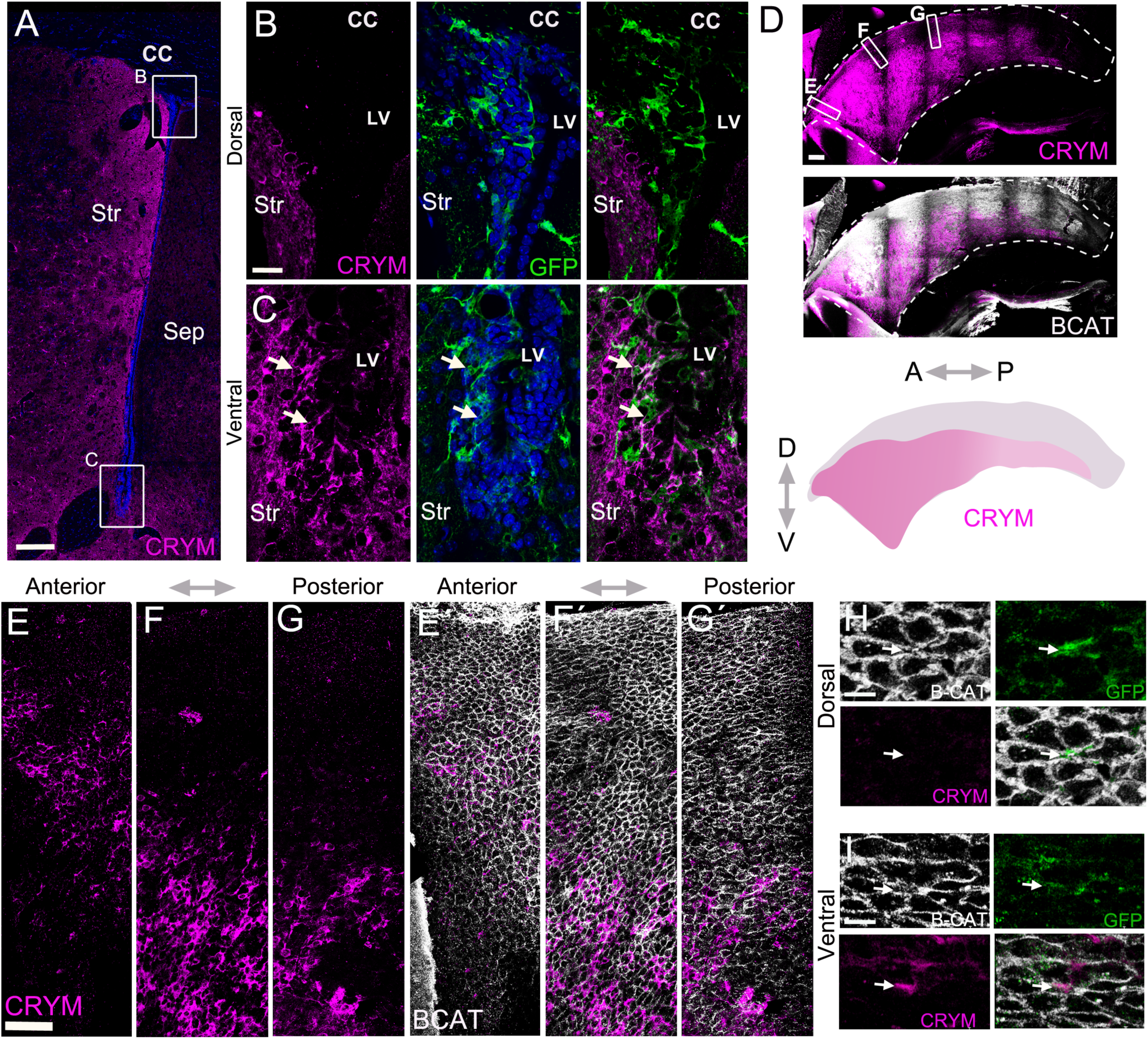
Whole mounts reveal CRYM expression in a wide ventral domain A. Confocal micrograph of a hGFAP:GFP coronal mouse brain section, where the V-SVZ is immunostained for GFP (green) and CRYM (magenta). B. High magnification image of the dorsal wedge region of the V-SVZ from (A). C. High magnification image of the ventral V-SVZ from (A). D. Immunostaining of CRYM (magenta) in a whole-mount preparation of the lateral wall of the V-SVZ, co-stained with 1-CATENIN (white), with a summary schematic depicting the extent of the CRYM+ domain. E. - G. Higher magnification images of boxed regions in (D) showing the distribution of CRYM+ (magenta) in V-SVZ cells outlined by 1-CATENIN (white). H. - I. High magnification images of GFP+ (green) B1 cell-containing pinwheels in the dorsal (H) and ventral (I) V-SVZ outlined by 1-CATENIN (white), also immunostained for CRYM (magenta). A: anterior, P: posterior, D: dorsal, V: ventral. Scale bars: 150 µm (A), 20 µm (B and C), 200 µm (D), 50 µm (E, F and G) and 10 µm (H and I).

### A cell cluster heterogeneity is linked to regionally-organized dorsal and ventral domains

We found that A cells (Fig 5A) were separated into two main sets of transcriptionally-related clusters: clusters A(15) and A(6) corresponded to A cells with a strong expression of genes associated with mitosis and cell cycle regulation (such as *Pclaf, Hmgb2, Rrm2, Mki67*, *Top2a*, and the *Mcm* gene family) (Fig 5B, Supplementary Table 3). We calculated the ‘area under the curve’ (AUC) scores (Aibar et al., 2017) for sets of genes corresponding to key GO terms in each A cell. The combined expression of genes in both the GO categories mitotic DNA replication (GO:1902969) and mitosis DNA replication initiation (GO:1902975) are highly upregulated in clusters A(15) and A(6) (Fig. 5-S1A, Supplementary Table 3). These clusters are also enriched in cells in S and G2M phases (Fig. 2K), indicating that these clusters correspond to dividing neuroblasts/early A cells (Lois and Alvarez-Buylla, 1993; Menezes et al., 1995). Clusters A(1), A(0), and A(4) corresponded to a second set of A cells expressing high levels of genes involved in cell migration, such as *Dab1* and *Slit2* (Fig. 5B; Supplementary Table 3). We found that neuron migration (GO:0001764) and spontaneous synaptic transmission (GO: 0098814) categories were more strongly present in clusters A(0), A(1) and A(4) (Fig. 5-S1A), indicating that these clusters likely correspond to migrating young neurons. This analysis of the GO terms enriched in A cells is consistent with the pseudotime analysis (Fig. 2L), indicating that A cells are organized in a continuum of maturation in UMAP space, with dividing neuroblasts at the top of the A cell cluster group and migrating young neurons in the bottom.

**Figure 5.**
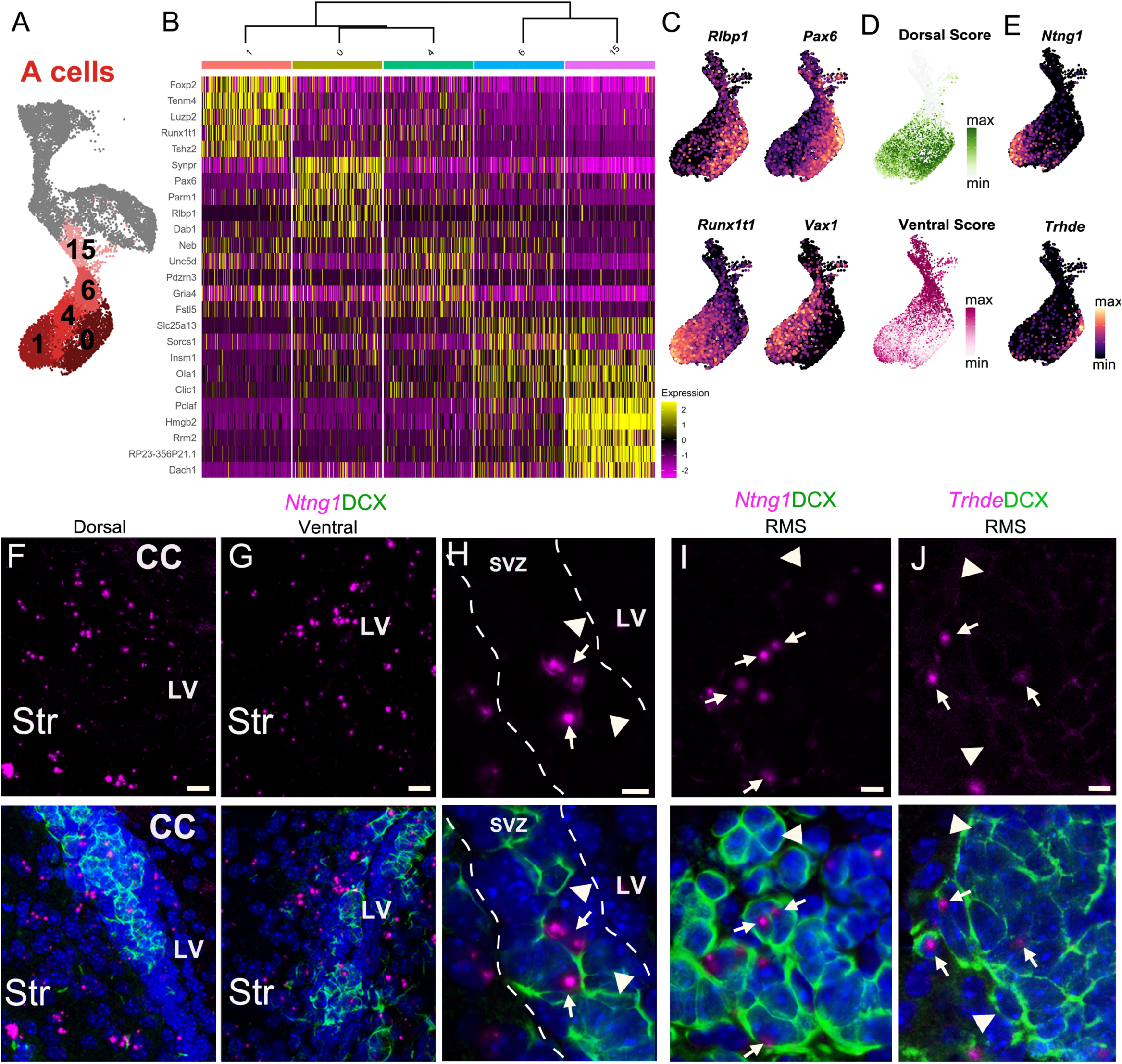
A cell cluster heterogeneity is linked to regionally-organized dorsal/ventral domains A. A cells are organized in 5 distinct clusters (1, 0, 4, 6, and 15). B. Heatmap showing the top 5 differentially expressed genes for each of the 5 clusters. C. The expression patterns of *Rlbp1* and *Runx1t1* are similar to *Vax1* and *Pax6*, respectively, which are genes known to be enriched in ventral and dorsal domains of the V-SVZ. D. The predicted region scores of each scRNA-Seq A cell plotted in UMAP space, where strongly Dorsal predicted identity cells are dark green, and strongly Ventral predicted identities are dark magenta. E. Expression patterns of marker genes *Ntng1* and *Trhde* in A cells. F. - H. Expression of *Ntng1* RNA (magenta) and DCX protein (green) in the dorsal (F) and ventral (G) V-SVZ, and high magnification of *Ntng1*-positive A cells (arrows) and *Ntng1*-negative A cells (arrowheads) in the V-SVZ (dotted lines). I. High magnification of *Ntng1* puncta in DCX+ A cells (arrows), and *Ntng1*-negative A cells (arrowheads) in the RMS. J. High magnification image of *Thrde* RNA (magenta) in DCX+ A cells (arrows), along with *Thrde*-negative DCX+ A cells (arrowheads) in the RMS. DAPI: blue, RMS: rostral migratory stream, LV: lateral ventricle, CC: corpus callosum, Str: striatum. Scale bars: 10 µm (F and G), 5 µm (H, I and J).

Interestingly, the combined expression of genes in dorsoventral axonal guidance (GO:0033563) and cerebral cortex regionalization (GO:0021796), terms associated with the regional specification of the brain, was high in clusters A(1) and A(0), despite individual GO terms not being statistically enriched in these clusters (Fig. 5-S1A; Supplementary Table 3). Additionally, *Runx1t1,* a transcription factor expressed in young neurons from the medial ganglionic eminence (Chen et al., 2017), and *Nxph1,* expressed in young migrating neurons from subpallial germinal zones (Batista-Brito et al., 2008), were among the most differentially expressed genes in cluster A(1) (Fig. 5B-C, Supplementary Table 3). Conversely, *Pax6,* a transcription factor that is highly expressed in the pallium and in the dorsal lateral ganglionic eminence (Ypsilanti and Rubenstein, 2016), was the most differentially expressed gene for cluster A(0) (Fig. 5B-C, Supplementary Table 3). Importantly, *Vax1* and *Pax6,* which have been previously found to be differentially expressed by ventrally- and dorsally-born A cells (Core et al., 2020), were highly expressed in clusters A(1) and A(0), respectively (Fig 5B-C). Taken together, gene expression suggests that A cells in cluster A(1) originate from ventral progenitors and A cells in cluster A(0) originate from dorsal progenitors. To test this hypothesis, we took advantage of the regional microdissections from the sNucRNA-Seq dataset and calculated dorsal and ventral predicted location scores for each scRNA-Seq A cell (Fig 5D). Intriguingly, among the more mature A cell clusters, we found that cluster A(0) was enriched in highly dorsal-scoring cells with relatively few highly ventral-scoring cells, while cluster A(1) was enriched in highly ventral-scoring cells with some highly dorsal-scoring cells (Fig 5D, Fig 5-S1D).

In order to confirm that A cells in A(0) and A(1) correspond to dorsal and ventral young neurons, respectively, we looked for markers of A(1) and A(0) that were minimally present in the other cluster. *Trhde* and *Ntng1* are expressed in subpopulations of A cells with high dorsal and ventral scores, respectively and we used RNAscope *in situ* hybridization with DCX immunolabeling to visualize the localization of transcripts in A cells *in vivo*. As suggested by the expression pattern in the scRNA-Seq clusters, *Ntng1* is not uniformly expressed in all A cells, but rather a subset of them (Fig 5F-I), in both dorsal and ventral V-SVZ (Fig 5F-H). *Trhde* puncta were present in the V-SVZ, along the ventricular lining and in areas immediately lateral to the DAPI-dense band of V-SVZ B, C and A cells (Fig 5-S1B). This expression pattern is consistent with *Trhde* expression in both the ependymal and striatal neuron scRNA-Seq clusters (Fig 5-S1C). We found few *Trhde+* DCX+ cells, in either the dorsal or ventral V-SVZ, where each positive A cell had one or two puncta that did not co-localize with the nucleus (Fig 5-S1B). In addition to *Ntng1 and Trhde* being found in DCX+ A cells dispersed along the full dorso-ventral axis of the V-SVZ (Fig 5F-H; Fig 5-S1B), in the DCX-dense RMS corridor, we found relatively small subsets of A cells that were *Ntng1+* or *Trhde*+, with only one or two mRNA puncta per cell (Fig 5I-J). Given the intermixing of A cells as they undergo tangential chain migration in the V-SVZ, the dorso-ventral sites of A cell origins cannot be visualized and validated by immunostaining or RNAscope (Fig 5 F-H, Fig 5-S1B). This is consistent with the expected spatial migration patterns of ventrally- vs. dorsally-born A cells (Fiorelli et al., 2015). Overall, we found that gene expression patterns, as well as independent, unbiased cell identity prediction provided additional support to the hypothesis that *Pax6^high^;Rlbp1^high^* cluster A(0) represents primarily dorsally-born A cells, while *Runx1t1^high^;Vax1^high^* cluster A(1) represents primarily ventrally-born A cells (Fig 5B-D). Together these data suggest that A cells are a molecularly heterogeneous population, and that heterogeneity is indicative of an A cell’s region of origin within the V-SVZ.

### Specific markers link regionally distinct B cell and A cell lineages

Interestingly, A(1) marker *Slit2* (Supplementary Table 3) and A(0) marker *Pax6* (Fig 5B-C) were also highly expressed in clusters corresponding to ventral and dorsal B cells, respectively (Fig. 3I; Supplementary Table 2; see below). In addition, *Rlbp1*, one of the main markers of dorsal B cells (Fig 3I), was also strongly expressed in cluster A(0) (Fig. 5C). To further determine if each dorsal/ventral domain has a specific gene expression signature that persists throughout the neurogenic lineage, we asked which B and A cell subpopulation marker genes were commonly expressed between the ventral B and ventral A cell clusters (B(14) and A(1)), and which were commonly expressed between the dorsal B(5+22) and dorsal A(0) clusters (Fig 6A-B). We identified 4 genes that were expressed throughout the dorsal lineage (*Rlbp1*, *Gm29260*, *Pax6*, and *Dcc*) and 5 genes expressed through the ventral lineage (*Adgrl3*, *Slit2*, *Ptprn2*, *Rbms1*, *Sntb1*) (Fig 6C). The converse comparison, however, of ventral B(14) and dorsal A(0), and dorsal B(5+22) and ventral A(1) clusters, yielded only one or two potential lineage marker genes, respectively (Fig 6-S1A-B). This suggests that B(14) and A(1), and B(5+22) and A(0) had a higher degree of transcriptional overlap, and correspond to ventral and dorsal lineages, respectively.

**Figure 6.**
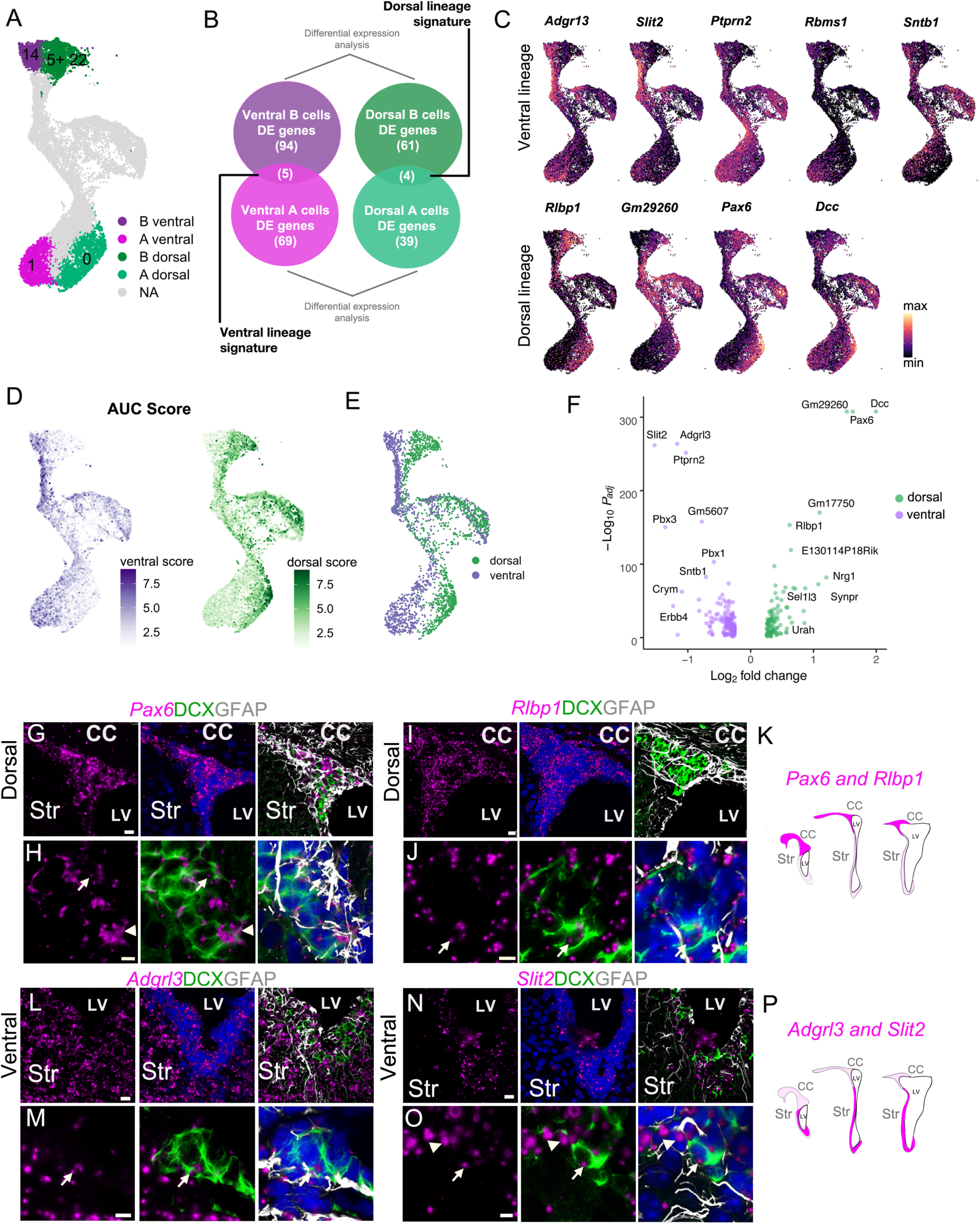
Transcriptional signatures of adult NSC regional heterogeneity are manifested along the neurogenic lineage A. UMAP plot highlighting the putative dorsal and ventral B and A cell clusters. B. Schematic illustrating the approach to identify genes that are differentially enriched in dorsal B and A, and ventral B and A cells, comparing B(14) to A(1) and B(5) & B(22) to A(0). C. Expression patterns of ventral (top row) and dorsal markers (bottom row) identified as differentially enriched throughout the B-C-A lineage. D. AUCell scoring of ventral (purple) and dorsal (green) cells based on the combined expression of genes through the entire B-C-A lineage. E. Classification of dorsal (green) and ventral (purple) cells in the neurogenic lineage based on the top quartile of AUCell scores. F. Volcano plot of significantly differentially expressed genes between the entire dorsal and ventral neurogenic lineages. G. - H. RNAscope validation of dorsal lineage marker *Pax6* (magenta) with DCX (green) and GFAP (white) immunostaining. High-magnification images of the V-SVZ dorsal wedge (G). H. High-magnification image of the ventral V-SVZ where *Pax6* colocalizes with an A cell (arrow) and B cell (arrowhead). I. - J. RNAscope validation of dorsal lineage marker *Rlbp1* (magenta) in the dorsal wedge (I). J. High-magnification images of *Rlbp1* in the dorsal V-SVZ, where puncta are visible in a DCX-positive A cell (arrow). K. Schematic illustrating the extent and expression levels of dorsal makers *Pax6* and *Rlbp1* across anterior-to-posterior sections of the V-SVZ. L. - M. RNAscope validation of ventral lineage marker *Adgrl3* (magenta) in the ventral V-SVZ. M. High-magnification image of *Adgrl3* in the ventral V-SVZ, colocalizing with an A cell (arrow). N. - O. RNAscope validation of ventral lineage marker *Slit2* (magenta) in the ventral V-SVZ. O. High-magnification image of the ventral V-SVZ, where *Slit2* puncta colocalize with an A cell (arrow) and B cell (arrowhead). P. Schematic illustrating the extent and expression levels of ventral makers *Adgrl3* and *Slit2* across anterior-to-posterior sections of the V-SVZ. CC: corpus callosum, Str: striatum, LV: lateral ventricle. Scale bars: 15 µm (G, L and N), 10 µm (I) and 5 µm (H, J, M and O).

To understand what other molecular differences exist between the putative dorsal and ventral lineages identified using only B and A cell cluster marker genes, we used the regional gene sets we identified above to calculate a composite AUC score (Aibar et al., 2017) for both the dorsal and ventral gene expression signatures (Fig 6D, Fig 6-S1C). We found that most cells scored highly for either the dorsal or ventral gene set, with highly ventral scoring cells located on the left side of the ‘bird’, and highly dorsal-scoring cells on the right (Fig 6D, Fig 6-S1C-E). When we plotted dorsal and ventral scores along pseudotime, we found that average dorsal and ventral scores were relatively constant along the B-C-A cell lineage progression, further suggesting these gene sets are stably expressed by dorsal and ventral cells during adult neurogenesis (Fig 6-S1F).

We then asked what other genes varied between the dorsal and ventral neurogenic lineages. By taking the top 25% of dorsal- and ventral-scoring cells (Fig. 6E, Fig 6-S1D-F), we identified 257 significantly differentially expressed genes between the dorsal and ventral neurogenic lineages: 108 dorsal markers and 149 ventral markers (Fig. 6F; Supplementary Table 4). We then conducted GO analysis on markers of the dorsal and ventral lineages. Ventral markers were enriched for genes associated with neurite outgrowth and migratory programs, including chemotaxis (GO:0050919), axon guidance (GO:0008045, GO:0031290), and V-SVZ-to-OB migration (GO:0022028). Dorsal markers were enriched for genes associated with glial development (GO:0021781), cortex regionalization (GO:0021796), and copper ion response (GO:0010273). Both dorsal and ventral marker sets were enriched for genes associated with OB interneuron differentiation (GO:0021889) (Fig. 6-S1G; Supplementary Table 4).

To validate RNA expression of putative dorsal and ventral lineage markers *in vivo,* we combined RNAscope labeling with GFAP and DCX immunostaining in coronal sections of the V-SVZ. We found that the putative dorsal lineage genes *Pax6* and *Rlbp1* were highly enriched in the dorsal region of the V-SVZ, with particularly enriched expression in the “wedge” region (Fig. 6G-K; Fig 6-S1H-K). Conversely, the putative ventral markers *Adgrl3* and *Slit2* had a higher expression in the ventral domain of the V-SVZ (Fig. 6L-P; Fig 6-S1L-O). Consistent with our scRNA-Seq data, these genes were expressed in both B and A cells in the neurogenic lineage (Fig 6H, J, M, O).

## Discussion

During brain development, regional allocations of the neuroepithelium give NSCs different neurogenic properties. The adult V-SVZ neurogenic niche retains regionally specified NSCs that generate different subtypes of neurons destined for the OB. A molecular understanding of what makes adult NSCs different between regions is largely lacking. Our scRNA-Seq and sNucRNA-Seq datasets provide new information about the diverse cell types that populate the V-SVZ. Beyond what previous studies have shown, our lineage analysis reveals parallel pathways of neurogenesis initiated by different populations of B cells. Interestingly, these differences in B cell identity correlate with unique regional patterns of gene expression, which we validated using reference-based metadata label transfer from a second dataset of regionally-dissected single V-SVZ nuclei. We confirmed the regional expression of marker genes by immunostaining and RNAscope analysis.

Our dataset provides sets of genes that are differentially expressed in dorsal and ventral B cells. Among these genes, we found well known regionally expressed transcription factors such as *Pax6*, *Hopx*, *Nkx6.2*, *Gsx2,* and *Vax1* (Core et al., 2020; Delgado and Lim, 2015; Hack et al., 2005; Kohwi et al., 2005; Merkle et al., 2014; Taglialatela et al., 2004; Zweifel et al., 2018). We also identified *Urah, Dio2* and *Crym* as novel markers that define largely non-overlapping domains of the V-SVZ. *Urah* and *Dio2* define a dorsal domain that includes the wedge and subcallosal roof of the V-SVZ, and *Crym* defines a wide ventral domain. Consistent with the above observations, *Crym* expression has been described in a subpopulation of qNSC in the early postnatal V-SVZ derived from the Nkx2.1 domain (Borrett et al., 2020). Our dorsal domain overlaps with that defined by *Pax6* (Hack et al., 2005; Kohwi et al., 2005) and the ventral domain with that defined by *Vax1* (Core et al., 2020). *Pax6* and *Vax1* are transcription factors that link these territories to well defined embryonic domains involved with the generation of different subsets of neurons in the cortex and striatum. Similarly, *Gsx2* is expressed in a gradient in the embryo, with its highest expression in the dorsal lateral ganglionic eminence (Corbin et al., 2000; Lopez-Juarez et al., 2013; Taglialatela et al., 2004; Waclaw et al., 2009; Young et al., 2007). Consistent with the dorsal expression pattern, in our dataset *Gsx2* was highly enriched in cluster B(5) (Fig. 3B). The dorsal region was also enriched in *Ptprz1*, *Hopx*, *Dio2*, *Tnc*, and *Moxd1,* which are also markers of outer radial glia, a subpopulation of human neural stem cells that continue to generate neurons for the cortex after detaching from the pallial wall of the lateral ventricles during prenatal development (Nowakowski et al., 2016; Pollen et al., 2015). These links with developmentally and regionally regulated genes will help define how the adult neurogenic niche inherits its regional specified domains.

Regional differences between ventral and dorsal domains of the V-SVZ had previously been inferred based on the labeling of non-overlapping territories using regional viral lineage tracing (Merkle et al., 2014, 2007; Ventura and Goldman, 2007). This analysis revealed different subtypes of OB interneurons derived from different NSC territories. Lineage tracing from embryonic to adult stages indicates that regional specification of neural stem cells is established early in embryonic development (Fuentealba et al., 2015). Genetic labeling of the most ventral domain of the V-SVZ showed the specific contribution of Nkx2.1-expressing B cells to deep layer granule cell neurons in the OB (Delgado and Lim, 2015). The ventral gene expression program is maintained by an epigenetic mechanism across cell divisions. In the absence of myeloid/lymphoid or mixed-lineage leukemia protein 1 (MLL1)-dependent epigenetic maintenance, the neurogenic lineage shifts to produce aberrantly ‘dorsalized’ OB neuronal subtypes (Delgado et al., 2020). This underscores the importance of regional identity in determining V-SVZ neuronal output. In our analysis, we also found region-associated gene expression programs contributing to A cell identity, along with a subset of dorsal- and ventral-specific genes expressed through the entire putative dorsal and ventral neurogenic lineages. Regional genes maintained through the neurogenic lineage could help us understand how NSC identities are maintained to ensure the production of the right types of interneurons in the OB.

The border between the *Crym*+ and the *Urah/Dio2*+ territories had not been previously determined and lies in an anatomically undefined location (Fig 4). Interestingly, the dorsal domain defined by *Dio2* and *Urah* largely overlaps with a region of high *Gli1* expression during neonatal and early postnatal stages (Tong et al., 2015). This dorsal Sonic Hedgehog-regulated domain in early postnatal life has been linked to oligodendrogenesis. Whether the adult domain we now unravel is developmentally linked to this early oligodendrogenic domain remains to be determined using lineage analysis. However, it is tempting to speculate that these territories are functionally linked, a hypothesis supported by the enrichment of glial-development-associated genes among dorsal markers (Fig 6G).

As indicated above, our analysis shows that *Crym, Urah,* and *Dio2* are differentially expressed by B cells according to region. *Crym, Urah,* and *Dio2* are all associated with the thyroid hormone signaling pathway; *Dio2* catalyzes thyroid hormone activation, *Crym* has been described to bind thyroid hormone, and *Urah* belongs to a family of thyroid hormone transporters (Luongo et al., 2019; Rudqvist et al., 2012; Vie et al., 1997). Thyroid hormone signaling has been shown to regulate V-SVZ neurogenesis (Lemkine et al., 2005; Lopez-Juarez et al., 2012; Luongo et al., 2021). Our findings suggest that hormone signaling may differentially affect B cells in different regions of the V-SVZ and therefore modify the balance of neuronal types produced for the OB. Region-specific regulation of V-SVZ stem cells has been previously demonstrated; hypothalamic projections innervate the anterior-ventral V-SVZ and activate Nkx2.1+ stem cells (Paul et al., 2017). The combination of region-specific innervation and molecular composition of B cells could form the basis for a system of hormonal regulation underlying circuit changes in the OB.

In summary, we present a large-scale single-cell description of dorso-ventral identity in the lateral wall of the V-SVZ. Not only do we recapitulate known divisions between dorsal and ventral B cells, but we identify novel regional B cell markers and uncover gene expression programs that appear to persist throughout lineage transitions (Fig 7). These data form a basis for future investigation of NSC identity and lineage commitment, providing clues to help us understand how molecularly-defined stem cell territories are spatially organized, and what distinguishes V-SVZ regions from one another.

**Figure 7.**
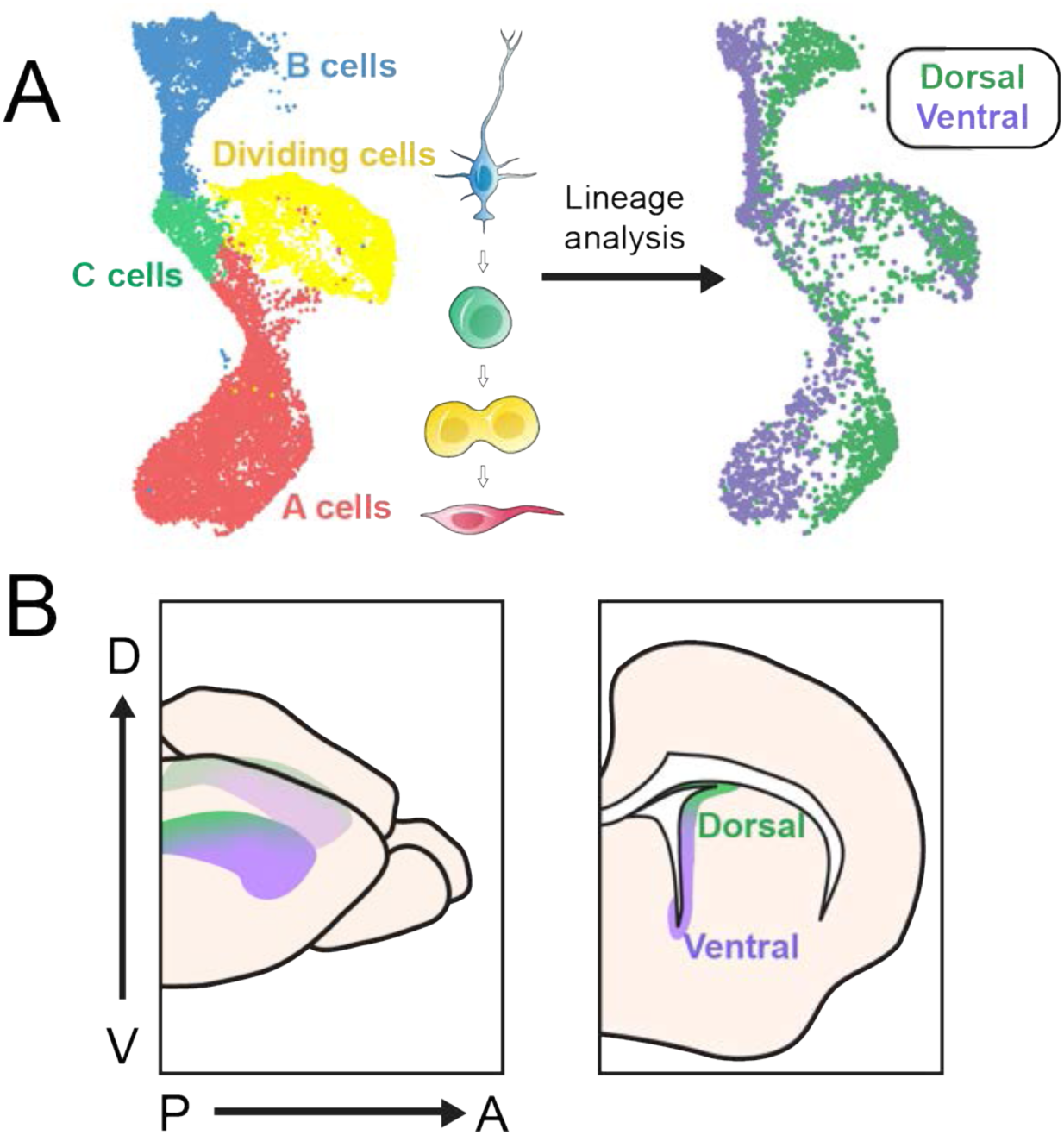
scRNA-Seq reveals dorsal and ventral neurogenic lineage domains in the V-SVZ A. A summary of cell types in the neurogenic lineage identified by scRNA-Seq and their classification into dorsal and ventral transcriptional identities. B. Schematic depicting the dorsal and ventral domains newly identified by scRNA-Seq and snucRNA-Seq, and confirmed by staining and RNAscope.

## Materials and Methods

### Mice

Mice were housed on a 12h day-night cycle with free access to water and food in a specific pathogen-free facility in social cages (up to 5 mice/cage) and treated according to the guidelines from the UCSF. Institutional Animal Care and Use Committee (IACUC) and NIH. All mice used in this study were healthy and immuno-competent, and did not undergo previous procedures unrelated to the experiment. CD1-elite mice (Charles River Laboratories) and hGFAP::GFP (FVB/N-Tg(GFAPGFP)14Mes/J, The Jackson Laboratory (003257)) (Zhuo et al., 1997) lines were used.

### Single Whole Cell Sample Preparation and Multiplexing

Mice received intraperitoneal administration of 2.5% Avertin followed by decapitation. Brains were extracted and 1 mm slices were obtained with an adult mouse brain slicer (Steel Brain Matrix - Coronal 0.5mm, Alto). Four samples were processed: sample 1: 2 males P35; sample 2: 2 males P35; sample 3: 2 females P29; and sample 4: 2 females P29. The lateral ventricle walls were microdissected in L-15 medium on ice and the tissue was transferred to Papain-EBSS (LK003150, Worthington). Tissue was digested for 30 mins at 37°C in a thermomixer at 900 RPM. Mechanical dissociation with a P1000 pipette tip (20 seconds), then fire-polished pasteur pipette was performed for 5 min. Tissue was digested for 10 more min at 37°C, and dissociated with the pasteur pipette for another 2 min. Cells were centrifuged for 5min, 300 RCF at room temp, and the pellet was resuspended with DNAase/ovomucoid inhibitor according to manufacturer’s protocol (Worthington). Cells were incubated in Red blood cell lysis buffer (420301, Biolegend) 3-4 min at 4°C. For MULTI-seq barcoding, cells were suspended with Anchor:Barcode solution (every sample was labeled with a unique barcode: sample 1 Barcode: TGAGACCT (“A3”); sample 2 barcode GCACACGC (“A4”); sample 3 barcode AGAGAGAG (“A5”); and sample 4 barcode TCACAGCA (“A6”)) for 5 minutes at 4°C. A Co-Anchor solution was added and incubated for 5 minutes (McGinnis et al., 2019). Samples were combined and filtered with a FlowMi 40µm filter (BAH136800040-50EA, Sigma). To remove myelin the cell suspension was incubated with Myelin Removal Beads (130-096-733, Miltenyi Biotec) (6µl/brain) for 15 mins at 2-8°C. Cells were washed with 0.5% BSA-PBS and transferred to MACS columns (30-042-401 and QuadroMACS Separator 130-090-976, Miltenyi Biotec). The cell suspension was preincubated with TruStain FcX Plus Antibody (BioLegend, Supplementary Table 5) on ice for 10 minutes, then incubated with oligonucleotide-tagged anti-VCAM1 and anti-CD24 antibodies (BioLegend, Supplementary Table 5) on ice for 30 minutes, then washed twice with 0.5% BSA-PBS by centrifugation (5 min, 4°C, 350 RCF) and filtered with a FlowMi 40µm filter. The effluent was collected and cell density was counted. Cells were loaded into two wells of a 10x Genomics Chromium Single Cell Controller. We used the 10x Genomics Chromium Single Cell 3’ Library & Gel Bead Kit v3 to generate cDNA libraries for sequencing according to manufacturer’s protocols. GFP expression of isolated cells was observed under an epifluorescence microscope.

### MULTI-seq Barcode Library Preparation & Cell Assignment

MULTI-seq and antibody TotalSeq barcode libraries were assembled as previously described (McGinnis et al., 2019). Briefly, a MULTI-seq primer is added to the cDNA amplification mix. Afterwards, in the first clean-up step using SPRI beads (0.6x) of the standard 10x library prep workflow, the supernatant is saved, transferred to a new tube and a cleanup step using SPRI (1.6x) is performed to eliminate larger molecules. A library preparation PCR is also performed for the MULTI-seq barcodes. The barcode library is analyzed using a Bioanalyzer High Sensitivity DNA system and then sequenced. The code for demultiplexing samples and detecting doublets can be found at https://github.com/chris-mcginnis-ucsf/MULTI-seq.

### Whole Cell Sequencing Data Alignment & Processing

We pooled gene expression and barcode cDNA libraries from each 10x Genomics Single Cell Controller well (technical replicates, “Lane”) and sequenced them at the UCSF Center for Advanced Technology on one lane of an Illumina Novaseq 6000 machine. A total of 2,892,555,503 reads were aligned using CellRanger 3.0.2-v3.2.0 (10x Genomics) to a custom version of the mouse reference genome GRCm38 that included the GFP gene (GFP sequence: Supplementary Table 6). Reads corresponding to oligonucleotide-tagged TotalSeq antibodies were assigned to cells in CellRanger according to manufacturer instructions.

To identify cell barcodes that most likely corresponded to viable cells, we performed quality control and filtering steps. We excluded cells outside of the following thresholds: UMI count depth: 5th and 95th percentiles; number of genes per cell: below 5th percentile); percentage of mitochondrial gene reads per cell: greater than 10%. We classified cells into sample groups and identified doublets using MULTI-seq barcode abundances (McGinnis et al., 2019). We used Seurat Integration (Seurat 3) canonical correlation analysis (CCA) to reduce data dimensionality and align the data from technical replicates (Lane 1 and Lane 2) (Stuart et al., 2019).

### Single Nucleus Sample Preparation

Brains were extracted and 0.5mm slices were obtained. We microdissected the anterior ventral, anterior-dorsal, posterior-ventral and dorsal V-SVZ regions of 17 P35 CD1 male (8) and female (9) mice. Briefly, we used a brain matrix to cut one millimeter thick coronal slabs of the mouse forebrain and used histological landmarks to identify each sampling area (e.g. anterior region landmarks: septum; posterior regions: hippocampus). Regions were dissected under a microscope to reduce the amount of underlying striatum in each sample. Each micro-dissected V-SVZ region was processed in parallel as a distinct sample. We processed tissue samples for nucleus isolation and sNucRNA-Seq as previously described (Velmeshev et al., 2019). Briefly, we generated a single nucleus suspension using a tissue douncer (Thomas Scientific, Cat # 3431D76) in nucleus isolation medium (0.32M sucrose, 5 mM CaCl2, 3 mM MgAc2, 0.1 mM EDTA, 10 mM Tris-HCl, 1 mM DTT, 0.1% Triton X-100 in DEPC-treated water). Debris was removed via ultracentrifugation on a sucrose cushion (1.8M sucrose, 3 mM MgAc2, 1 mM DTT, 10 mM Tris-HCl in DEPC-treated water) in a thick-walled ultracentrifuge tube (Beckman Coulter, Cat # 355631) and spun at 107,000 RCF, 4°C for 150min. The pelleted nuclei were incubated in 250µL PBS made with DEPC-treated water on ice for 20 minutes. The resuspended pellet was filtered twice through a 30µm cell strainer. We counted nuclei with a hemocytometer to determine nucleus density, and loaded approximately 12,000 nuclei from each sample into its own well/lane of a 10x Genomics Chromium Single Cell Controller microfluidics instrument. We used the 10x Genomics Chromium Single Cell 3’ Library & Gel Bead Kit v2 to generate cDNA libraries for sequencing according to manufacturers’ protocols. We measured cDNA library fragment size and concentration with a Bioanalyzer (Agilent Genomics).

### Single Nucleus Sequencing Data Alignment & Processing

We pooled the gene expression cDNA libraries from each single nucleus sample and sequenced them on one lane of an Illumina HiSeq 4000 at the UCSF Center for Advanced Technology. The PV sample was further sequenced to increase sequencing depth. A total of 1,340,031,643 reads were aligned using CellRanger 2.1.0-2.3.0 (AV, AD, PD samples); 3.0.2-v3.2.0 (PV sample) (10x Genomics) to a custom mouse reference genome that includes unspliced “pre-mRNA” (GRCm38), which we expect to be present in cell nuclei (Velmeshev et al., 2019).

To identify cell barcodes that most likely corresponded to viable nuclei, we performed quality control and filtering steps. For each region sample, we excluded nuclei outside of the following thresholds: UMI count depth: 5th to 95th percentiles; number of genes per cell: below 5th percentile; fraction of mitochondrial gene reads per cell (<10%). We used Seurat Integration (Seurat 3) canonical correlation analysis (CCA) to reduce data dimensionality and align the data from each region (Stuart et al., 2019).

### Single Cell & Single Nucleus Sequencing Data Normalization and Dimensionality Reduction

We used Seurat 3 (Stuart et al. 2019) to analyze both the Whole Cell and Single Nucleus datasets: for each dataset, cells or nuclei from each 10x Chromium Controller Lane (scRNA-Seq: Lanes 1 and 2; sNucRNA-Seq: AD, AV, PD, PV lanes) were integrated using IntegrateData and normalized using regularized negative binomial regression (SCTransform) (Hafemeister and Satija, 2019). We calculated 100 principal components (PCs) per dataset, and used 50 (scRNA-Seq) or 100 (sNucRNA-Seq) to calculate cell cluster identities at five distinct resolutions (0.5, 0.8, 1.0, 1.5 and 2.0) and UMAP coordinates. The cell cluster identities presented in this manuscript correspond to resolution 1.5 (scRNA-Seq metadata column integrated_snn_res.1.5) or 2 (sNucRNA-Seq metadata column integrated_snn_res.2), and were chosen based on visual correspondence with the expression of known neurogenic lineage markers. Sequenced antibody tags in the scRNA-Seq dataset were separately normalized using NormalizeData (method: CLR) and ScaleData, and are included in the Seurat object ‘Protein’ assay as “VCAM1-TotalA” and “CD24-TotalA”.

### Dual Fluorescent *in situ* Hybridization-Immunofluoresce

Mouse brains (n=2, P30) were serially sectioned using a Leica cryostat (10 um-thick sections in Superfrost Plus slides). Sections were incubated 10 min with 4% PFA and washed 3×10 min with phosphate-buffered saline (PBS) to remove OCT. Slides were incubated with ACD hydrogen peroxide for 10 min, treated in 1x target retrieval buffer (ACD) for 5 min (at 96-100 °C) and rinsed in water and 100% ethanol. Samples were air dried at 60°C during 15 min and kept at room temperature overnight. The day after, samples were treated with Protease Plus for 30 min at 40 °C in the RNAscope oven. Hybridization of probes and amplification solutions was performed according to the manufacturer’s instructions. Amplification and detection steps were performed using the RNAscope 2.5 HD Red Detection Kit (ACD, 320497) and RNAscope 2.5 HD Duplex Reagent Kit (ACD, 322430). RNAscope probes used: Mm-Lphn3 (also named Adgrl3) (cat.# 317481), Mm-Rlbp1 (cat.# 468161), Mm-Crym (cat.# 466131), Mm-Pax6 (cat.# 412821), Mm-Slit2 (cat.# 449691), Mm-Cnatnp2 (cat.# 449381), Mm-Urah-C2 (cat.# 525331-C2), Mm-Dio2 (cat.# 479331), Mm-Hopx (cat.# 405161), Mm-Ntng1-C2 (cat.# 488871-C2), Mm-Trhde (cat.# 450781). Mm-Snhg15 was custom made (NPR-0009896, cat.# 889191). DapB mRNA probe (cat.# 310043) was used as negative and Mm-PPIB (cat.# 313911) as positive control. RNAscope assay was directly followed by antibody staining for chicken anti-GFAP (Abcam, ab4674,1:500) and rabbit anti-DCX (Cell signaling, 4604S, 1:200) or rabbit anti-S100 (Dako, Z033, 1:100, *discontinued*) (Supplementary Table 5). Samples were blocked with TNB solution (0.1 M Tris-HCl, pH 7.5, 0.15 M NaCl, 0.5% PerkinElmer TSA blocking reagent) 30 min and incubated in primary antibodies overnight. Samples were washed with PBS-Tx0.1% and incubated with secondary antibodies Donkey anti-Chicken Alexa 647 (Jackson ImmunoResearch, 703-605-155, 1:500) and Donkey anti-Rabbit biotinylated (Jackson ImmunoResearch, 711-065-152, 1:400) in TNB buffer for 1.5 hrs. Samples were washed and incubated with Streptavidin HRP (1:200 in TNB solution) for 30 min. Washed 3×5 min and incubated with Fluorescein Tyramide 5 min (1:50 in amplification diluent) rinsed and incubated with DAPI 10 min. Sections were mounted with Prolong glass Antifade Mountant (Invitrogen, P36980).

### Immunohistochemistry

Coronal sections (n=2 mice, P30) and whole mounts (n=2 mice, P28) were incubated with Immunosaver (1:200; EMS, Fort Washington, PA) for 20 min at 60°C, and then 15 min at RT. Tissue was then incubated in blocking solution (10% donkey serum and 0.2% Triton X-100 in 0.1 M PBS) for 1hr. followed by overnight incubation at 4°DC with the primary antibodies: mouse anti-CRYM (Santa Cruz, sc-376687,1:100), rabbit anti-BETA-CATENIN (Sigma, C2206, 1:250), Chicken anti-GFP (Aves labs, GFP1020, 1:400), Rabbit anti-HOPX (Proteintech,11419-1-AP,1:500)(Supplementary Table 5). On the next day, sections were rinsed and incubated with Alexa Fluor secondary antibodies. Samples were mounted with Aqua-poly/mount (Polysciences Inc, 18606-20).

### Confocal Microscopy

Confocal images were acquired using the Leica Sp8 confocal microscope. Samples processed for RNAscope and immunohistochemistry were imaged at 20x (low magnification) and 63x (High magnification). For high magnification RNAscope images, 10-15 optical sections were acquired sequentially and analyzed using Leica Application Suite X (LAS X) software.

### Differential Gene Expression and GO Analysis

We used Seurat 3 functions FindMarkers (two groups) or FindAllMarkers (more than two groups) to identify differentially expressed genes among groups of single cells. For detailed parameters see available code (below). Selection of genes from the resulting lists for further analysis are described in the text. Gene Ontology analyses of differentially expressed genes were performed in Panther v.16 (Mi et al., 2021). GO terms with a false discovery rate lower than 5% were considered statistically significant.

### Nuclei to Whole Cell Regional Label Transfer

B cell label transfer: First, we subsetted quiescent B cells from both the scRNA-Seq and sNucRNA-Seq datasets (clusters B(5), B(14), B(22), and sNucRNA-Seq cluster 7). We generated the reference sNucRNA-Seq B cell dataset that consisted of equal numbers of B cells per region, randomly selected from the middle 50% of cells by number of genes identified per cell (25th-75th percentile of SCT_snn_nFeature). This prevented the region with the most nuclei from dominating the prediction scores, and filtering cells by nFeature prior to downsampling resulted in reproducible prediction scoring, likely due to exclusion of low quality B cells and doublets not rejected in the full dataset quality control steps. Anterior dorsal and posterior dorsal regions were combined to create the Dorsal reference cell set, and the anterior ventral and posterior ventral were combined to create the Ventral reference cell set. Subsetted scRNA-Seq B cells and filtered sNucRNA-Seq B cell sets were individually normalized using SCTransform. We then ran FindTransferAnchors with the following settings: *reference.assay = “SCT”, query.assay = “SCT”, normalization.method = “SCT”, npcs=30, project.query=T, and dims = 1:30*. We then calculated Dorsal and Ventral predicted identity scores for each scRNA-Seq B cell using TransferData (Stuart et al., 2019).

A cell label transfer: The same method as above was applied to scRNA-Seq A cell clusters A(0), A(1), A(4), A(6), and A(15), and sNucRNA-Seq clusters 12 and 29.

### RNA Velocity

RNA Velocity in the neurogenic lineage was calculated using scvelo (Bergen et al., 2020), using 2000 genes per cell. Moments were calculated using 30 PCs and 30 neighbors. Velocity was estimated using the stochastic model. Pseudotime was plotted using the original UMAP coordinates.

### Data & Code Availability

The RNA sequencing datasets generated for this manuscript are deposited in the following locations: scRNA-Seq and sNucRNA-Seq GEO Data Series: <u>GSE165555</u>.

Processed data (CellRanger output .mtx and .tsv files, and Seurat Object .rds files) are available as supplementary files within the scRNA-Seq (<u>GSE165554</u>) or sNucRNA-Seq (<u>GSE165551</u>) data series or individual sample entries listed within each data series.

Web-based, interactive versions of the scRNA-Seq and sNucRNA-Seq datasets are available from the University of California Santa Cruz Cell Browser: https://svzneurogeniclineage.cells.ucsc.edu

The code used to analyze the datasets and generate the figures are available at the following location: https://github.com/AlvarezBuyllaLab?tab=repositories

## Acknowledgments

We would like to thank Christopher McGinnis and Dr. Zev Gartner for the MULTI-seq barcodes and technical advice. We thank Dr. Aparna Bhaduri, Dr. Alex Pollen and Dr. Dmitry Velmeshev for input on single cell experimental design and analysis. This work was supported by grants from the US National Institutes of Health R37 HD032116, R01 NS028478, R01 NS113910, a generous gift from the John G. Bowes Research Fund and the UCSF Program for Breakthrough Biomedical Research, partially funded by the Sandler Foundation (to A.A.B); R01 NS091544, R01 NS112357 and VA 1I01 BX000252 (to D.A.L.); F32 NS103221 (to S.A.R.); F31 NS106868 (to D.W.); and by the Spanish Generalitat Valenciana and European Social Fund (APOSTD2018/A113) (to A.C.S). A.A.B. is the Heather and Melanie Muss Endowed Chair and Professor of Neurological Surgery at UCSF. A.A.B. is Cofounder and on the Scientific Advisory Board of Neurona Therapeutics.

---

**Figure 1 - Supplement 1.**
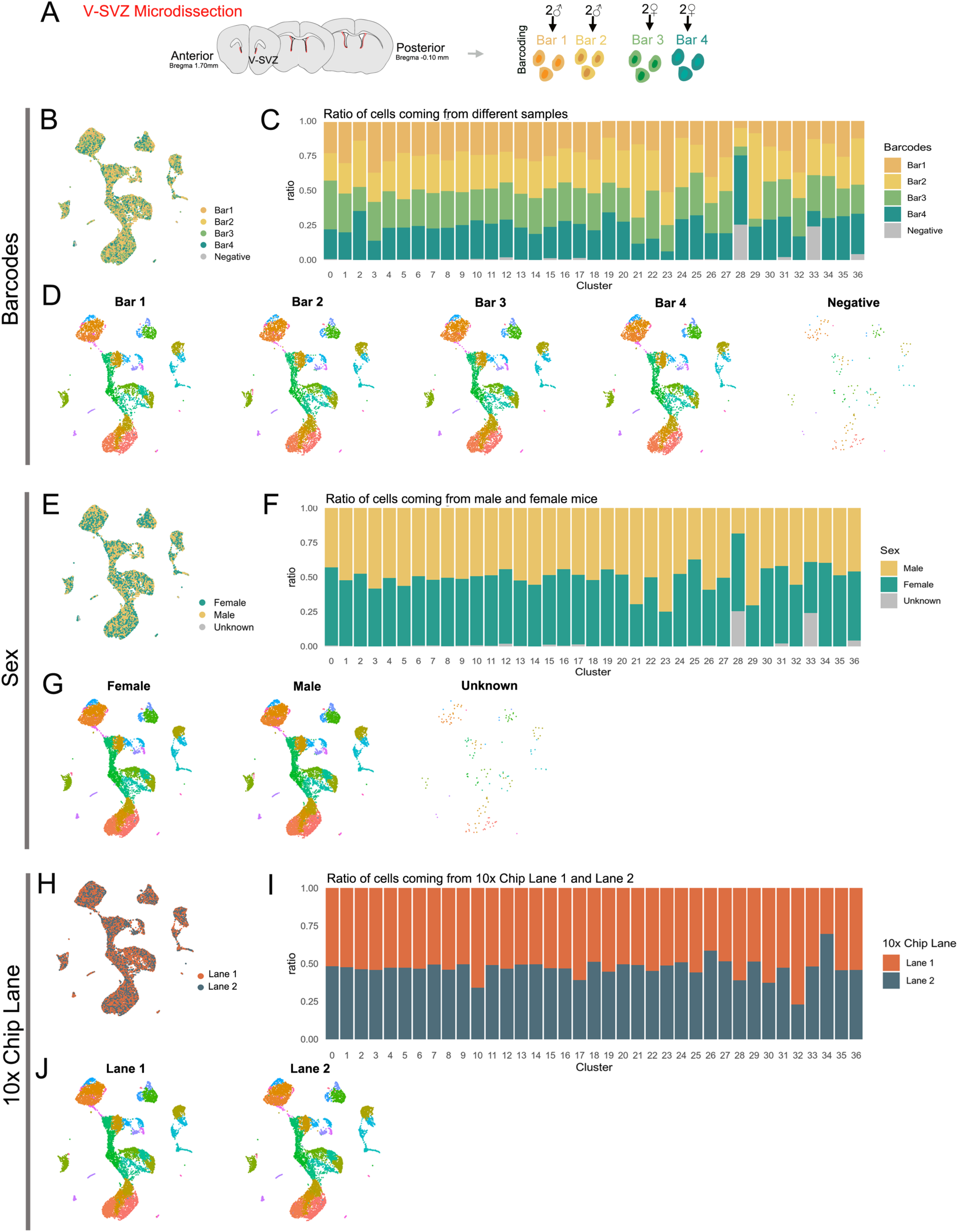
Biological and technical replicate metadata of scRNA-Seq dataset A. Diagram representing V-SVZ microdissected areas (red). The V-SVZ was microdissected from 1mm brain slices from anterior (bregma 1.70) to Posterior bregma (-0.10 mm) regions. Single cell suspensions from two female samples and two male samples were multiplexed with MULTI-seq barcodes (Bar1-4). B. UMAP plot of sequenced cells after demultiplexing and quality control filtering, color-coded by MULTI-seq barcode. Cells with no detectable barcode are labeled “Negative”. C. Ratio of cells within each cluster (see Fig 1A, right panel) coming from each Barcode sample. D. UMAP plots of cells separated by Barcode number and color-coded by cluster number (see Fig 1A, right panel). E. UMAP plot of sequenced cells color-coded by animal sex: Female cells (green) correspond to Barcodes 3 & 4 combined, male cells (tan) correspond to Barcodes 1 & 2 combined. Cells with no detectable barcode are labeled “Unknown” (gray). F. Ratio of cells within each cluster (see Fig 1A, right panel) coming from male (tan) or female (green) mice, or of unknown origin (gray). G. UMAP plots of cells separated by animal sex and color-coded by cluster number (see Fig 1A, right panel). H. UMAP plot of sequenced cells color-coded by 10x Chromium Controller Chip lane (equivalent to “batch”, or technical replicate): Lane 1 (orange) or Lane 2 (navy). I. Ratio of cells within each cluster (see Fig 1A, right panel) coming from Lane 1 (orange) or Lane 2 (navy). J. UMAP plots of cells separated by Lane and color-coded by cluster number (see Fig 1A, right panel).

**Figure 1 - Supplement 2.**
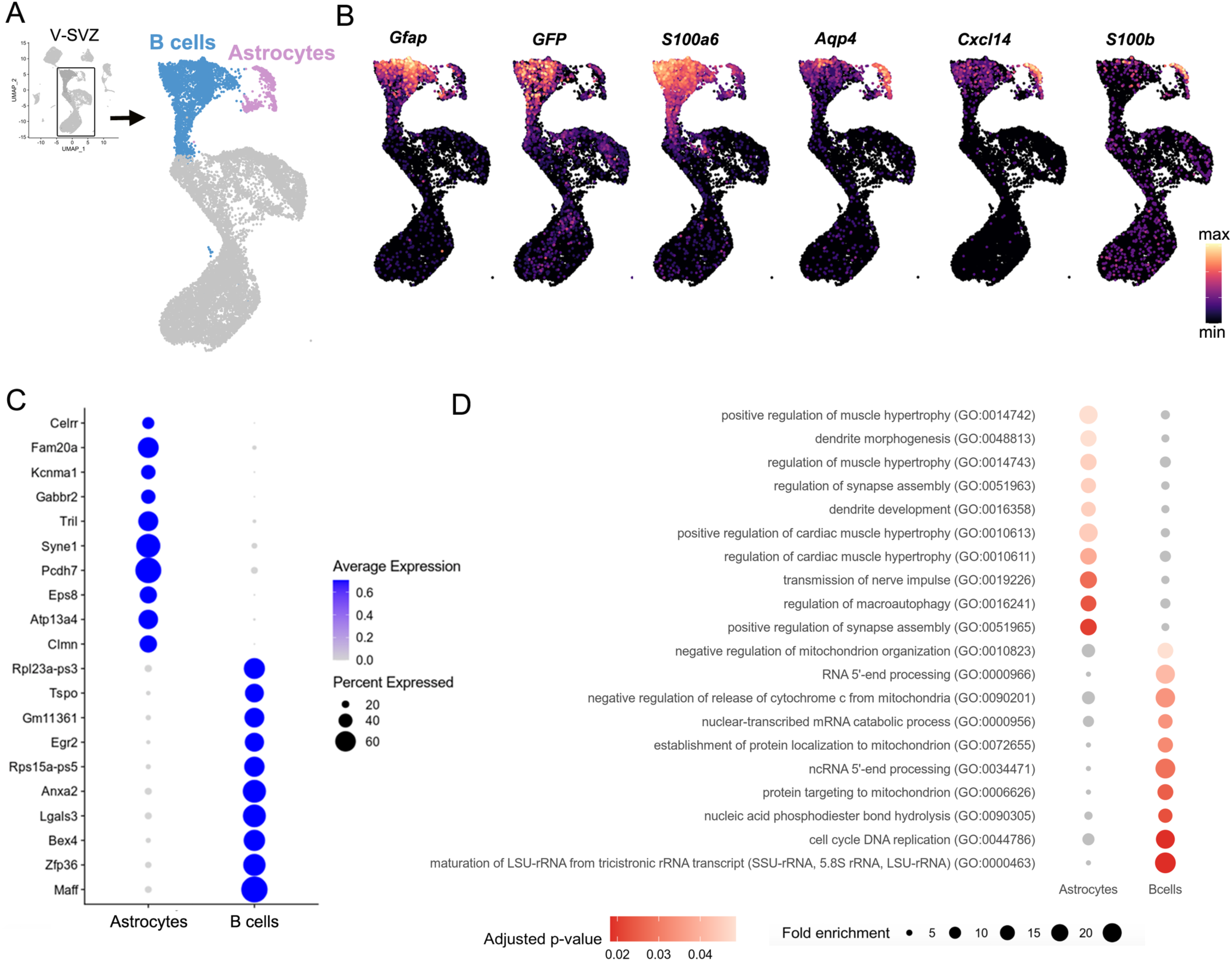
Transcriptomic profile of B cells versus Astrocytes A. UMAP plot of V-SVZ neurogenic lineage cells (blue) and astrocytes (purple) isolated from the scRNA-Seq dataset (inset, boxed area). B. Gene expression of B cell-enriched markers *Gfap, GFP, S100a6* (top row) and astrocyte-enriched markers *Aqp4, Cxcl14, S100b* (bottom row). C. Dot plot of the top ten differentially expressed Astrocyte and B cell cluster markers. D. Dot plot of significantly over-represented Gene Ontology (GO) terms identified by gene set enrichment analysis using differentially expressed Astrocyte and B cell genes.

**Figure 1 - Supplement 3.**
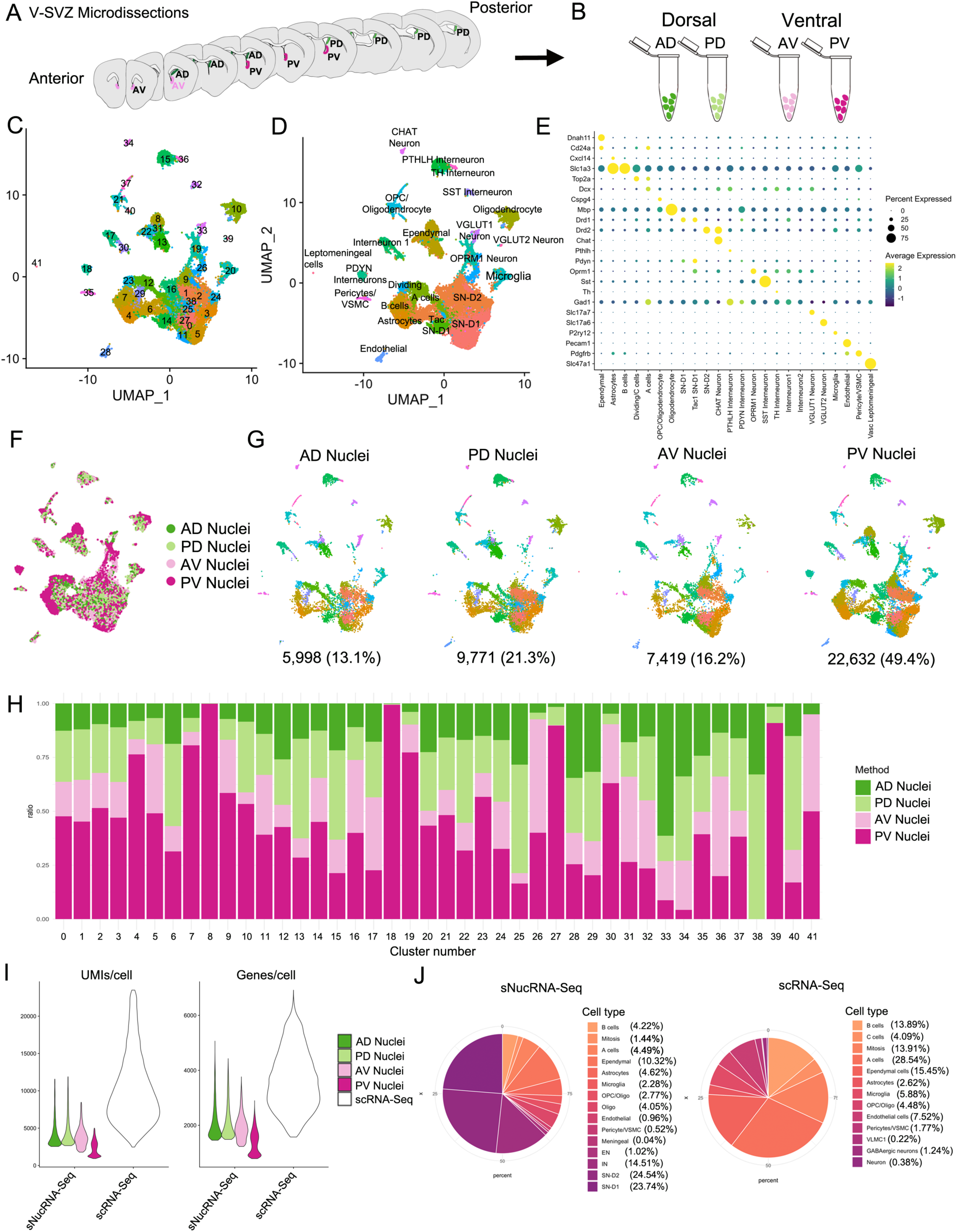
Characterization of the sNucRNA-Seq dataset A. Diagram representing the four V-SVZ microdissected regions: Anterior ventral (AV, pink), anterior dorsal (AD, dark green), posterior ventral (PV, magenta), and posterior dorsal (PD, light green). B. Single nucleus suspensions from each region were processed as separate samples in parallel. C. UMAP plot of sequenced nuclei after quality control filtering, color-coded by cluster number. D. UMAP plot of sNucRNA-Seq cell types. E. Dot plot of cell type specific marker expression in the clusters from (D). F. UMAP plot of sequenced cells color-coded by region: AD (green), PD (light green), AV (pink), and PV (magenta). G. UMAP plots of nuclei separated by region and color-coded by cluster number (see panel C). Below each plot is the number of nuclei and the percent of the total dataset originating from each region. H. Ratio of nuclei within each cluster (see panel C) coming from each region sample. I. Violin plot of the number of unique molecular identifiers (UMIs) detected per nucleus or cell (left panel), and number of genes detected per cell (right panel) in each sNucRNA-Seq region and in the scRNA-Seq dataset (white). Median UMIs: 9,181^scRNA-Seq^/3,062^sNucRNA-Seq^, a 3-fold difference. Median genes: 3,549^scRNA-Seq^/1,679^sNucRNA-Seq^, a 2.1-fold difference). J. Pie charts of cell types represented in sNucRNA-Seq (left) and scRNA-Seq datasets (right). The scRNA-Seq dataset contains 60.4% neurogenic lineage cells (shades of orange; 14,660/24,261 total cells), compared to 10.6% in the sNucRNA-Seq dataset (4,859/45,820 total nuclei). Conversely, the sNucRNA-Seq dataset contains 61.5% neurons (shades of purple; 28,185 nuclei, 13 subtypes (panel D), while the scRNA-Seq contains only 1.6% (395 cells, 3 subtypes (Fig 1A-B). The scRNA-Seq and sNucRNA-Seq datasets each contain 37.9% (9,206 cells) and 25.1% (11,480 nuclei) glial cells, respectively (shades of plum). Cell types listed in each pie chart legend are plotted in order, clockwise from coordinate 0.

**Figure 3 - Supplement 1.**
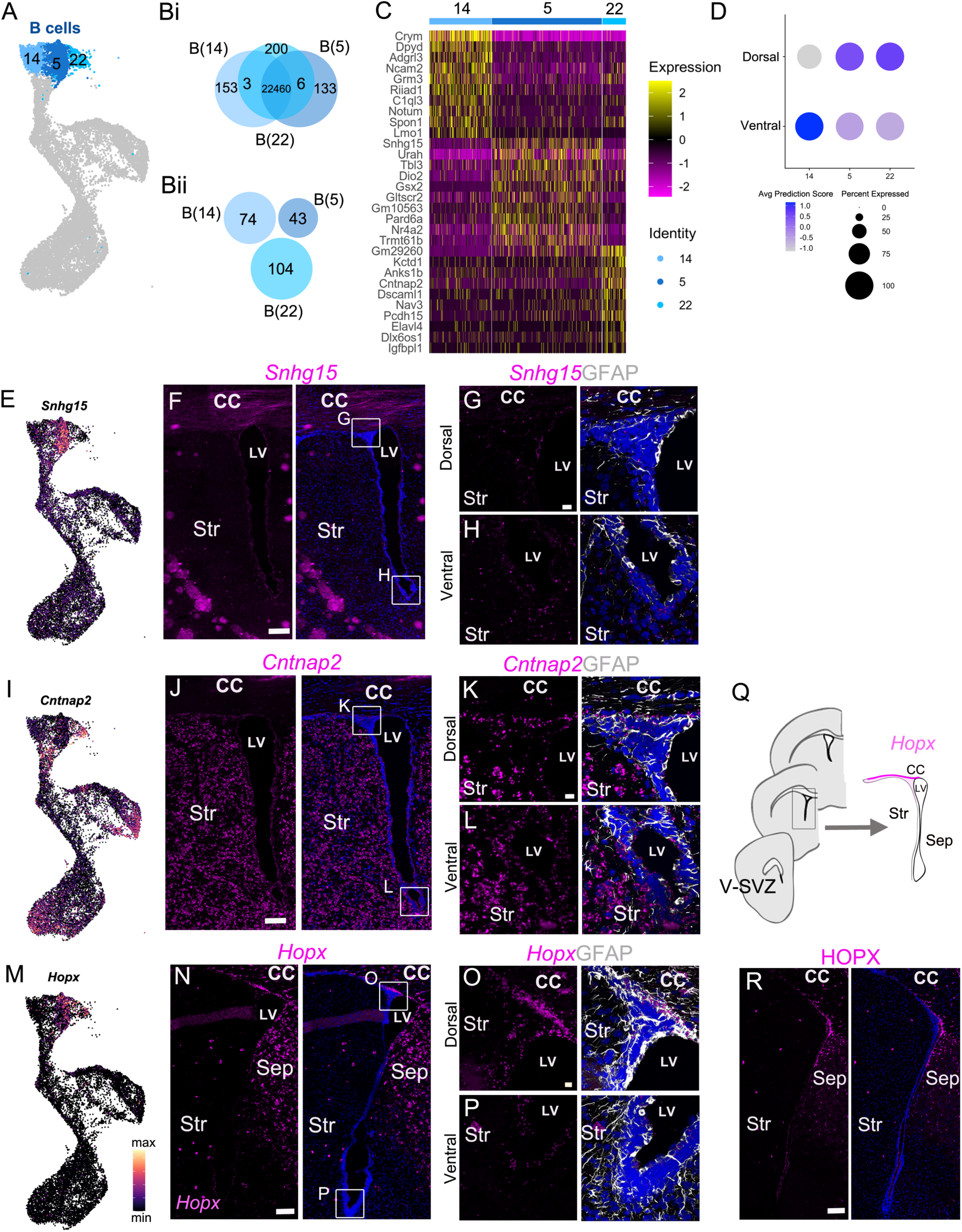
A. UMAP plot of B cell cluster identities used in the following analysis: B(14), B(5) and B(22). B. ***i.*** Venn diagram summarizing differential gene expression analysis between clusters B(14), B(5) and B(22). ***ii.*** Numbers of candidate marker genes identified after selecting significantly upregulated genes expressed in no more than 40% of cells of the other cluster. C. Heatmap depicting expression of the top 10 differentially expressed genes between clusters B(14) (left), B(5) (center) and B(22) (right). D. Dot plot of the average Dorsal or Ventral predicted identity scores for scRNA-Seq B cell clusters B(14), B(5) and B(22). E. UMAP plot of *Snhg15* expression in the scRNA-Seq neurogenic lineage. F. Confocal micrograph of *Snhg15* RNA (magenta) expression in the V-SVZ. Insets denote locations of high magnification images (G) and (H). G. - H. High magnification of dorsal (G) and ventral (H) V-SVZ regions denoted in (F) showing *Snhg15* RNA (magenta) and GFAP protein (white) expression. I. UMAP plot of *Cntnap2* expression in the scRNA-Seq neurogenic lineage. J. Confocal micrograph of *Cntnap2* RNA (magenta) expression in the V-SVZ. Insets denote locations of high magnification images (G) and (H). K. - L. High magnification of dorsal (K) and ventral (L) V-SVZ regions denoted in (J) showing *Cntnap2* RNA (magenta) and GFAP protein (white) expression. M. UMAP plot of *Hopx* expression in the scRNA-Seq neurogenic lineage. N. Confocal micrograph of *Hopx* RNA (magenta) expression in the V-SVZ. Insets denote locations of high magnification images (O) and (P). O. - P. High magnification of dorsal (O) and ventral (P) V-SVZ regions denoted in (N) showing *Cntnap2* RNA (magenta) and GFAP protein (white) expression. Q. Summary schematic of anterior to posterior coronal brain sections analyzed (left) and one of the V-SVZ regions (boxed area on left, enlarged on right). *Hopx* expression is summarized as strong (magenta), sparse (light magenta) or absent (white) in different subregions of the V-SVZ. R. Confocal micrograph of HOPX protein expression (magenta) in the V-SVZ. DAPI: blue, LV: lateral ventricle, CC: corpus callosum, Str: striatum. Scale bars: 100 µm (F, J, H and R) 10 µm (G, H, K, L, O and P).

**Figure 5 - Supplement 1.**
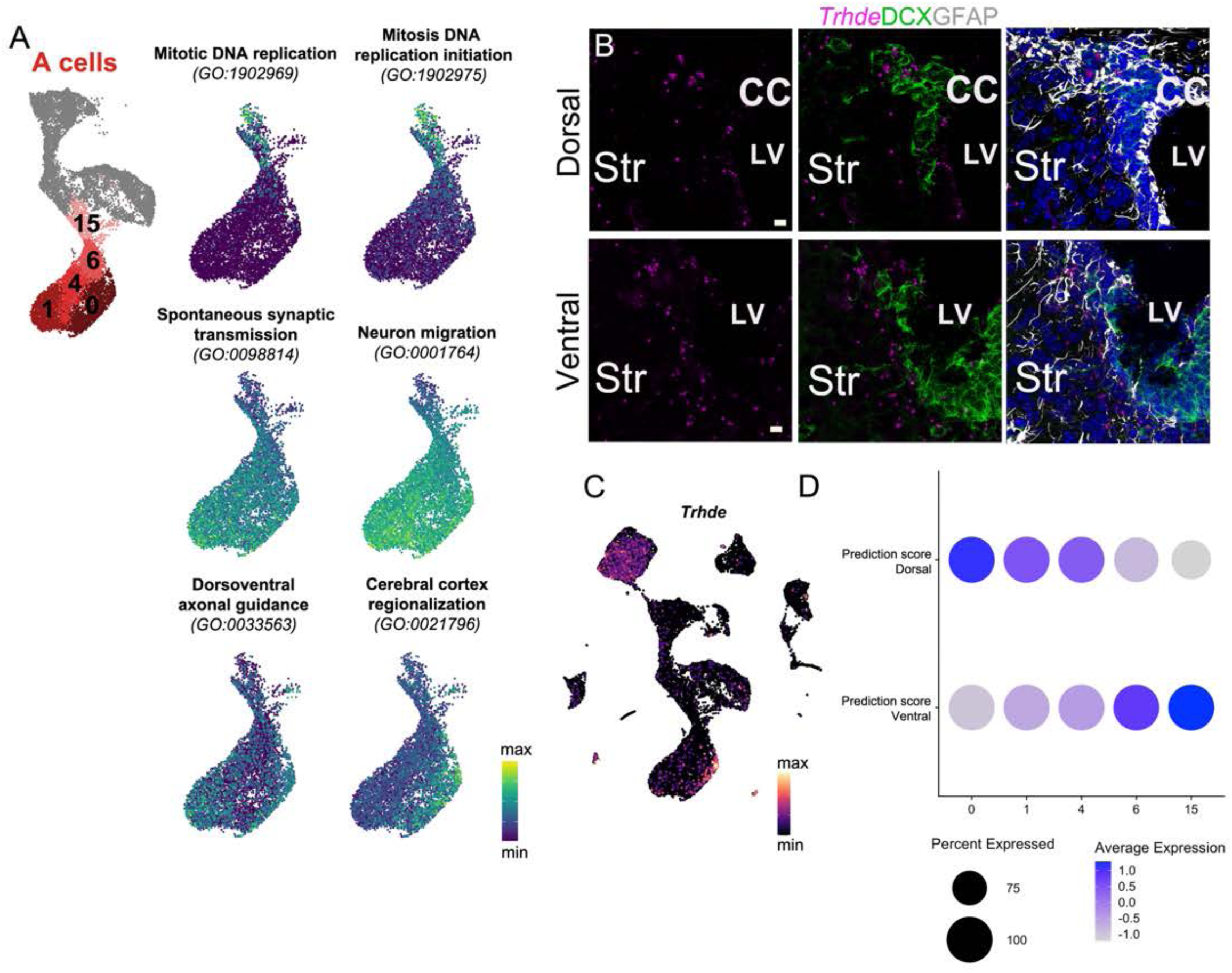
A. UMAP plot of A cell clusters, and UMAP plots of only A cell clusters labeled with AUCell scores of combined gene expression of genes contained in each of the six GO categories shown. B. *Trhde* expression (magenta) in the dorsal (top row) and ventral (bottom row) V-SVZ, colabled with DCX (green) and GFAP (white) immunostaining. C. *Trhde* expression in the full scRNA-Seq dataset. D. Dot plot of the average dorsal and ventral prediction scores for each of the A cell clusters 0, 1, 4, 6 and 15. DAPI: blue, LV: lateral ventricle, CC: corpus callosum, Str: striatum. Scale bar: 10 µm (B)

**Figure 6 - Supplement 1.**
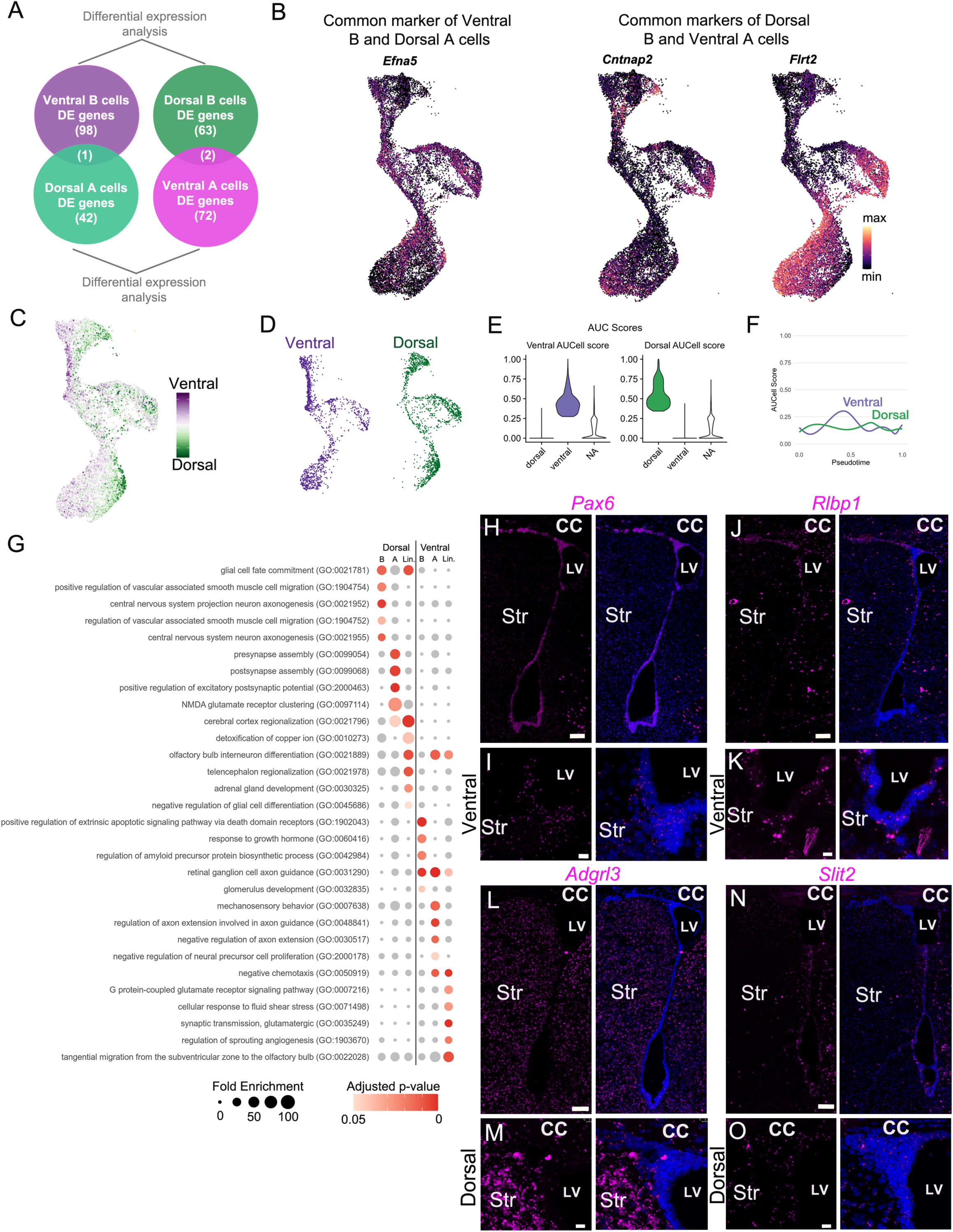
A. Schematic illustrating the control comparison to identify genes enriched in both ventral B and dorsal A cells, or in dorsal B and ventral A cells. B. UMAP plots showing expression of the 3 genes identified in the analysis in (A), including *Cntnap2*, *Flrt2*, and *Efna5*. C. UMAP plot colored by position score, based on net ventral (purple) and dorsal (green) AUCell scores. D. UMAP of the neurogenic lineage split by classification into the ventral (purple) or dorsal (green) lineage. E. Violin plot of AUCell score distribution of dorsal and ventral cells shown in (D). NA: cells not assigned. F. Ventral and dorsal AUCell scores plotted over pseudotime, as calculated by RNA velocity. G. Dot plot representing enriched GO terms identified by marker genes of the dorsal and ventral lineages (“Lin”), markers of dorsal vs. ventral B cells (“B”), or markers of dorsal vs. ventral A cells (“A”). H., J, L, N. Low-magnification images of the entire V-SVZ in a coronal section, labeled with RNAscope probes (magenta) against *Pax6* (H); *Rlbp1* (J); *Adgrl3* (L) and *Slit2* (N). I, K. High magnification of the ventral V-SVZ from panel H (I) and J (K). M, O. High magnification of the dorsal V-SVZ from panel L (M) and N (O). Scale bars: 100 µm (H, J, L and N) and 15 µm (I, K, M and O).

